# A robust model for cell type-specific interindividual variation in single-cell RNA sequencing data

**DOI:** 10.1101/2023.02.24.529987

**Authors:** Minhui Chen, Andy Dahl

## Abstract

The development of single-cell RNA sequencing (scRNA-seq) offers opportunities to characterize cellular heterogeneity at unprecedented resolution. Although scRNA-seq has been widely used to identify and characterize gene expression variation across cell types and cell states based on their average gene expression profiles, most studies ignore variation across individual donors. Modelling this inter-individual variation could improve statistical power to detect cell type-specific biology and inform the genes and cell types that underlying complex traits. We therefore develop a new model to detect and quantify cell type-specific variation across individuals called CTMM (Cell Type-specific linear Mixed Model). CTMM operates on cell type-specific pseudobulk expression and is fit with efficient methods that scale to hundreds of samples. We use extensive simulations to show that CTMM is powerful and unbiased in realistic settings. We also derive calibrated tests for cell type-specific interindividual variation, which is challenging given the modest sample sizes in scRNA-seq data. We apply CTMM to scRNA-seq data from human induced pluripotent stem cells to characterize the transcriptomic variation across donors as cells differentiate into endoderm. We find that almost 100% of transcriptome-wide variability between donors is differentiation stage-specific. CTMM also identifies individual genes with statistically significant stage-specific variability across samples, including 61 genes that do not have significant stage-specific mean expression. Finally, we extend CTMM to partition interindividual covariance between stages, which recapitulates the overall differentiation trajectory. Overall, CTMM is a powerful tool to characterize a novel dimension of cell type-specific biology in scRNA-seq.

## Introduction

The technology of single-cell RNA sequencing (scRNA-seq) profiles gene expression at the resolution of single cells. This resolution may be essential for understanding molecular mechanisms underlying many complex traits because disease gene expression is highly cell type-specific^1–3^. For example, *APOE* is a risk gene for Alzheimer’s disease that is downregulated in astrocytes but is upregulated in microglia^2^. One common application of scRNA-seq is to investigate differentially expressed genes (DEG) that exhibit differences in mean expression between cell types, such as diseased vs healthy^4^ or pre- vs post-treatment^5–7^. Furthermore, methods to infer cell type labels in scRNA-seq data primarily rely on differential mean expression between cell types^8,9^.

However, few scRNA-seq studies have evaluated gene expression variation across individuals. Understanding this variation could help identify and characterize genes and cell types that cause interindividual variation in complex traits ranging from height to autoimmune disorders. Studies using bulk RNA-seq have shown that gene expression variability informs disease biology and drug development^10–12^. However, bulk transcriptomics has poor resolution on individual cell types, which can cause both false positives and false negatives. In particular, prior signals in bulk RNA-seq could be explained by variation in cell type proportions rather than variation in gene expression within cell types^13^. Because scRNA-seq data has cell-level resolution, it provides an opportunity to powerfully partition expression variation within and between cell types. This has recently become possible with the proliferation of population-scale scRNA-seq studies that contain hundreds of individuals^14–18^.

In this paper, we develop CTMM (Cell Type-specific linear Mixed Model) to detect and quantify cell type-specific variation across individuals in scRNA-seq data. CTMM considers three nested models of cell type-shared and -specific variation. In the simplest model, all variation is shared homogeneously between cell types (“Hom”), with cell types differing only in mean expression. The next model allows independent variation in each cell type (“Free”), i.e., cell type-specific variation. The richest model allows for arbitrary forms of covariance between cell types (“Full”). We focus on fitting CTMM to cell type-specific pseudobulk, which averages gene expression over cells within each cell type for each individual. We also develop a version of CTMM for overall pseudobulk data, which averages over all cells from all cell types for each individual. Overall pseudobulk is less powerful and is akin to bulk sequencing data. We explored several statistical methods to fit and test CTMM’s parameters, which is nontrivial due to the modest sample sizes of scRNA-seq data. We performed a series of simulations to evaluate performance in a broad range of realistic settings. We then applied CTMM to characterize transcriptomic variation across individual donors along the developmental trajectory from human induced pluripotent stem cells (iPSCs) to endoderm. Transcriptome-wide, CTMM found that almost all interindividual variation was specific to each developmental time point, and the Full model found greater correlation between nearby time points. We also identified specific genes with statistically significant time point-specific variation across individuals, including genes with known importance for cell pluripotency and differentiation.

## Material and Methods

### Models

#### Overview of cell type-specific linear mixed models for gene expression

We model the expression level of a given gene for individual *i*, cell type *c,* and cell *s* by:

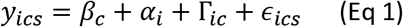

In this model, *y_ics_* is the gene expression level for the *s*-th cell from cell type *c* in individual *i*; note that the number of measured cells varies across individuals and cell types. *β_c_* is the mean expression level in cell type *c*, which we model as a fixed effect. *α_i_* captures differences between individuals that are shared across cell types, which we model as a random effect: 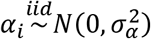. Γ_*ic*_ captures the difference between individuals that is specific to cell type *c*, which we also model as a random effect by 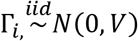. Here Γ_*i*_, is a vector of cell type-specific expression for individual *i* and *V* is a *C* × *C* matrix describing cell type-specific variances and covariances across cell types.

Finally, *ϵ_ics_* is the residual effect, which we assume to be *i.i.d.* for each individual-cell type pair with *E*_(*ϵ*_ics_)_ = 0 and 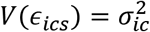. We directly estimate 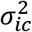 from the single cell-level data by 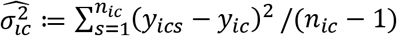, where *n_ic_* is the number of cells in individual *i* and cell type *c* and *y_ic_* is the average expression across all *n_ic_* cells (*i.e.,* cell type-specific pseudobulk, defined below). 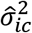. is unbiased even if *ϵ_ics_* is non-Gaussian, which is important because expression in single cells is non-Gaussian. Note that this is impossible in bulk expression data, even if sorted into cell types, because bulk only measures average expression. That is, scRNA-seq data makes it possible to distinguish true interindividual variation from measurement noise.

Our focus is the covariance matrix *V*, which captures the difference and similarity between cell types (for a given gene). The diagonal terms capture cell type-specific variance. If there is no cell type-specific variation between individuals, then *V_cc_* = 0 for all *c.* The off-diagonal terms capture covariance between specific pairs of cell types; if all cell types are equally similar to each other, then *V_cc′_* = 0 for all *c* ≠ *c′.*

We consider three nested models of interindividual variation defined by the structure of *V.* First, the homogeneous (Hom) model assumes that *V* = 0, *i.e.,* that all expression variance is shared homogeneously across cell types without any cell type-specificity. Second, the Free model allows arbitrary levels of cell type-specific variance by allowing *V* to be an arbitrary diagonal matrix. Third, the Full model captures arbitrary levels of covariance between specific cell type pairs by allowing *V* to be any positive semidefinite matrix. Intuitively, the Hom model captures variation across individuals, but assumes this variation is identically shared across cell types. The Free model allows cell type-specific variation, *e.g.,* a gene that is largely similar between individuals except in a single cell type. The Full models allows complex relationships among cell types, *e.g.*, hierarchical relationships among immune cell types.

A technical consideration in the Full model is that *V* and 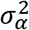 are not jointly identified. Specifically, passing a constant between 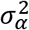 and *V* does not change the likelihood (*i.e.*, 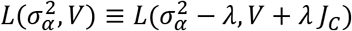, where *J_C_* is *C*×*C* matrix containing all 1s). Therefore, without loss of generality, we set 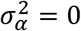 in the Full model. The Full model is statistically challenging because its number of parameters scales quadratically with the number of cell types, *C.* In practice, the Full model only has precise estimates with hundreds to thousands of samples or, as below, when aggregating together many genes.

#### Deriving models for overall and cell type-specific pseudobulk

Directly modelling single cell expression as in Eq1 is challenging computationally and statistically. Computationally, modelling individual cells increases the number of observations by orders of magnitude because there can be dozens or hundreds of cells per individual-cell type pair. Statistically, the individual cell’s expression is highly non-Gaussian, requiring additional assumptions and computationally expensive generalized linear mixed models. Instead, we study scRNA-seq data at the level of pseudobulk expression, which averages expression over many cells. We consider both overall pseudobulk (OP), which averages over all measured cells per individual, and cell type-specific pseudobulk (CTP), which averages over cells in each cell type per individual.

Specifically, the pseudobulk measures that we input to CTMM are:

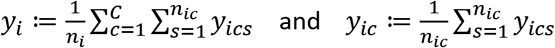

where *y_i_* is the OP expression for individual *i*, and *y_ic_* is the CTP expression for individual *í* and cell type *c*.

Our cell-level model in Eq 1 implies the following mixed model for the OP expression:

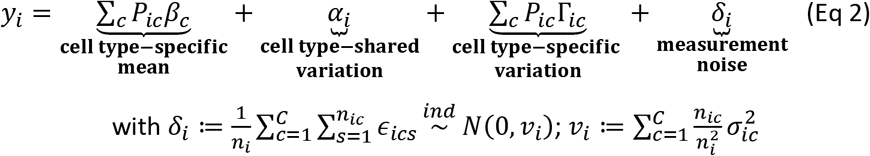

*δ_i_* is the measurement noise for individual *i*, with variance *v_i_* that we estimate by plugging in our estimate of 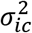. *P* is the matrix of cell type proportions, defined by 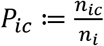.

Our cell-level model in Eq 1 also implies a mixed model on the CTP expression data:

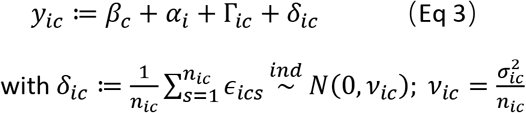

Here, *δ_ic_* is the noise for individual *i* and cell type *c*, with variance *v_ic_*. By the central limit theorem, both *δ_i_* and *δ_ic_* are approximately Gaussian when *n_ic_* is not too small, even though *ϵ_ics_* is very non-Gaussian.

### Fitting and testing parameters of CTMM

We evaluated three approaches to estimate the parameters in CTMM: maximum likelihood (ML), restricted maximum likelihood (REML), and method-of-moments (HE, as it is called Haseman-Elston regression in genetics). We implemented ML by maximizing the likelihood function using BFGS algorithm implemented in the R function ‘optim’ (Supplementary Note). REML was fit similarly using the restricted likelihood, which residualizes covariates from the full likelihood (Supplementary Note). For both REML and ML, we reran 10 random restarts if the initial optimization attempt failed (Supplementary Note), which is important to mitigate bias from local maxima with small sample sizes. We allowed negative variance components to reduce bias, though the total expression variance was always positive. Due to the complexity of these likelihood functions, we evaluated refining the BFGS solution with Nelder-Mead iterations; we found that this is not necessary for the analyses considered in the Main text, but it can be important for the more challenging analyses, such as fitting the Free model with ML on OP data (Supplementary Figure S1). We fit HE analytically (Supplementary Note).

Because the CTP expression data is a vector of length *NC*, where *N* is the number of individuals, naively fitting CTMM in ML and REML has a computational complexity of *O*(*N*^3^*C*^3^). We use several linear algebra identities to simplify the complexity to *O*(*NC*^3^). The relative gains will increase as *N* and *C* grow, which are both expected in future scRNA-seq datasets.

Our primary test compared the Hom and Free models, which asks whether interindividual variation is cell type-specific or shared uniformly across cell types. In other words, this is a test for differential expression variance across cell types. By comparison, standard tests for differential expression ask whether the mean expression levels, *β_c_*, differ across cell types. We also developed a test comparing the Full and Free models to ask whether cell types covary across individuals, but we found this has very low power at current sample sizes (data not shown).

We implemented likelihood ratio tests (LRT) and Wald tests to compare the Free and Hom models. For the LRT, we used *C* degrees of freedom because the Free model adds variance components for each cell type (but see Crainiceanu and Ruppert^19^ and Greven et al.^20^ for more detailed discussion of these tests). For the Wald test, we used an *F*-test with *C* numerator degrees of freedom and *N* – *R* denominator degrees of freedom, where *R* is the number of model parameters in the Free model, that is *2C* + 1. We evaluated two options to estimate the precision matrix for CTMM’s variance component estimates, which is needed for the Wald test. First, we used the inverse of the Fisher information matrix for REML and ML, which is consistent for large sample sizes. Second, we used jackknife (JK) to nonparametrically estimate the precision matrix by fitting the model after excluding each sample in turn. For large sample sizes, both LRT and Wald tests are valid; however, we are interested in modest sample sizes and hence we profile a wide range of approaches.

We tested for mean expression differentiation by evaluating the null hypothesis that *β_c_* = *β_c′_* for all cell types *c* and *c′. β* is the cell type fixed effect (*i.e.,* cell type-specific mean expression) and is estimated using generalized least squares with variance components fit under the Free model. We used a Wald test for *β* with numerator degrees of freedom *C* – 1 and denominator degrees of freedom *N* – *R*, where *R* is the number of parameters in the Free model, that is 2*C* + 1. We estimated the precision matrix for CTMM’s estimates of *β* using jackknife. The jackknife includes re-fitting variance components, which is important because these variance component estimates are noisy.

We have also extended CTMM to accommodate additional random effects (Supplementary Note). This can be essential in practice, but it can be computationally infeasible as it requires inverting large matrices. Fortunately, the primary use case involves blocked random effects, such as experimental batch in our iPSC analysis. We derived a different optimization approach designed specifically for this common scenario, which simplified the computational complexity by orders of magnitude. In our iPSC analysis, these manipulations reduced REML computation time per gene from ~40 seconds to ~10 second.

### Simulation

We tested the performance of CTMM with a series of simulations under Hom and Free models. We simulated gene expression for each individual from Eq 2 (for overall pseudobulk) and Eq 3 (for cell type-specific pseudobulk). We varied the number of individuals, cell type proportions, and levels of cell type-specific variance. For each simulated dataset, we evaluated all three methods to fit CTMM (ML, REML, and HE) and each applicable test for cell type-specific interindividual variance (LRT and Wald). For each simulation parameter setting, we ran 1,000 replicate simulations to calculate the average CTMM estimates, their sampling distributions, and the test positive rate. We also simulated under the more complex Full model. Further simulation details are provided in the Supplementary Note, with a list of simulation parameters in Supplementary Table S1.

We also performed simulations to assess CTMM’s sensitivity to estimation errors in *v_ic_*, the level of noise due to cell-level variation. This is important because *v_ic_* is not known in practice. Specifically, for each *v_ic_*, we draw *x_ic_ i.i.d.* from a *Beta*(2,*b*) distribution and then add +*x_ic_v_ic_* or –*x_ic_v_ic_* before inputting *v_ic_* to CTMM (Supplementary Note). To span the range of estimation errors in the real iPSCs data, we simulated *b* = 20, 10, 5, 3, 2. To evaluate power under a range of Free models, we varied cell type-specific variance for cell type 1 (*V*_11_) from 0.05 to 0.5 and fixed other cell type-specific variances to 0.1. For simplicity, the Free model simulations always used *b* = 5 (the most realistic value). As this simulation focuses on CTMM’s utility in our real data analysis, we simulated using the parameters we estimated in the iPSCs data below (Supplementary Note), and we only examined CTP as it is far more powerful. We ran 1,000 replicates for each setting of simulation parameters.

### Differentiating iPSCs analysis

#### Data and model

Human induced pluripotent stem cells (iPSCs) are derived from somatic cells that have been reprogrammed into an embryonic-like pluripotent state. iPSCs can differentiate to diverse cell types, with a concomitant transcriptomic trajectory across time as the cells differentiate. We studied the transcriptome as iPSCs differentiate into endoderm using scRNA-seq data from 125 individual donors^15^. Cells were collected on four consecutive days as the iPSCs differentiate, starting from iPSCs, which we used to define four cell types. We used the log transformed gene expression data provided by Cuomo *et al.,* which has been through a thorough process of quality control and normalization (https://zenodo.org/record/3625024#.Xil-0y2cZ0s). The dataset includes 11,231 genes and 36,044 cells.

For the 33 individuals who had technical replicates in the data, we only included the replicate with the largest number of cells. We excluded individuals with fewer than 100 cells to better satisfy the Gaussian approximation of *δ_ic_*, leaving 94 individuals.

For each gene, we fit OP and CTP expression into Hom, Free, and Full models with ML, REML, and HE. In all models, we adjusted for sex, neonatal diabetes, and the first 6 principal components calculated on OP expression as fixed effects. We used our extension of CTMM to model experimental batch as a random effect, which is important because batch has large effects that cannot be ignored yet has too many degrees of freedom to fit as fixed effects (24 batches vs 94 individuals). We used Bonferroni correction to account for multiple testing across genes.

#### Impute pseudobulk data

For individual-cell type pairs with no more than 10 cells, *y_ic_* and *v_ic_* were set to missing. We then imputed missing entries in *y_ic_* and *v_ic_*. We compared three approaches to imputation (each applied separately to *y* and *v*). First, we imputed each gene separately using either softImpute^21^ or MVN-impute (implemented in Dahl *et al.*^22^). In brief, the former makes a low-rank approximation, while the latter approximates individuals as independent and leverages correlations among cell types. We also evaluated imputing all genes jointly across the transcriptome using softImpute (in an *N* × *CG* matrix, where *G* is the number of genes); this is computationally impossible with MVN-impute.

To evaluate imputation accuracy, we masked observed entries in *y_ic_* and *v_ic_* and compared the imputed values to the masked true values. Of note, if one cell type of an individual has less than or equal to 10 cells, all genes’ expression would be missing for the pair of individual and cell type. To be realistic, we maintained this structure of missingness by employing a “copy-mask” approach as in our prior study^22^. We randomly sampled an individual with missing cell types and masked the same cell types in another randomly chosen individual. We repeated this process until 10% of all pairs of individual and cell type were masked. We calculated correlation and mean squared error (MSE) between imputed values and masked true values across individuals for each gene-cell type pair. We conducted 10 replications of the process of masking and imputation and calculated the medians of correlation and MSE across those repeats as final measures of imputation accuracy. For *v_ic_*, imputation might get negative values by chance. We treated these negative variances in different ways in OP and CTP expression data. In OP, we set negative variances to 0, so they had little impact on the estimation of *v_ic_* while maintaining information from other cell types; in CTP, for each gene and cell type, we set them to maximum raw *v_ic_* in that specific gene and cell type, so they contributed less to model likelihood. Note that standard approaches to impute expression in single cells^23,24^ does not impute the pseudobulk data, which has missing entries due to missing cells, not missing expression within observed cells.

## Results

### Simulation

We simulated a series of scenarios to assess the performance of CTMM. We simulated Hom and Free models by varying sample size, level of cell type-specific variance, and cell type proportions. We first evaluated the accuracy to quantify cell type-specific variance. Supplementary Figure S2 showed the estimation of cell type-specific variance in the simulation of Free model with varying sample sizes from 20 to 300. As expected, when fitting simulated data into the Free model, both OP and CTP performed well, as illustrated by the roughly unbiased estimates of cell type-specific variance *V.* The performance improved along with the increase of sample size. CTP provided more precise estimates than OP, since CTP uses more information than OP by modeling pseudobulk expression for each cell type. Comparing methods for parameter estimation, likelihood-based methods including ML and REML had similar level of precision, and both had better precision than HE, since likelihood-based methods utilize more information than HE by assuming normal distribution of random effects. Supplementary Figure S3 showed estimates with varying levels of cell type-specific variance. Our models provided unbiased estimates of cell type-specific variance, even in the simulation of Hom model where there is no cell type-specific effect. Figure S4 showed estimates with varying cell type proportions. When decreasing the proportion of main cell type (with the largest cell type-specific variance), all models performed well except for HE with CTP input, which broke down when the main cell type proportion went below 10%. We also simulated under the Full model, which had precise and unbiased estimates of covariance between cell types when sample size was above 50 with CTP (Supplementary Figure S5).

We then evaluated the power of our models to detect cell type-specific variance. Figure 1 showed positive rates of REML and HE using OP or CTP data as input for different sample sizes. Under the simulation of Hom model where there is no cell type-specific variance, different tests for cell type-specific variance with both OP and CTP input were appropriately null with around 5% of false positive rate, except for REML (Wald), that is Wald test in REML using precision matrix inferred from the Fisher information matrix. REML (JK), that is jackknife-based Wald test in REML, and HE were slightly inflated in CTP when sample size was 100 or lower. Under the simulation of Free model, CTP gained much larger power than OP, for example, when sample size was 50, CTP had 10-fold positive rate over OP (100% versus 10% using REML with LRT). REML (LRT) in CTP had the best power. Its true positive rate reached above 80% even when sample size was only 20 and reached 100% when sample size was 50. The other three tests in CTP, including REML (Wald), REML (JK), and HE, also had over 80% of true positive rate when sample size reached 100. ML and REML had similar performance when fitting CTP (Supplementary Figure S6). We also assessed the impact of cell type proportions and level of cell type-specific variance. As expected, the power decreased when the main cell type became rare and cell type-specific variance was low (Supplementary Figure S6).

**Figure 1.**
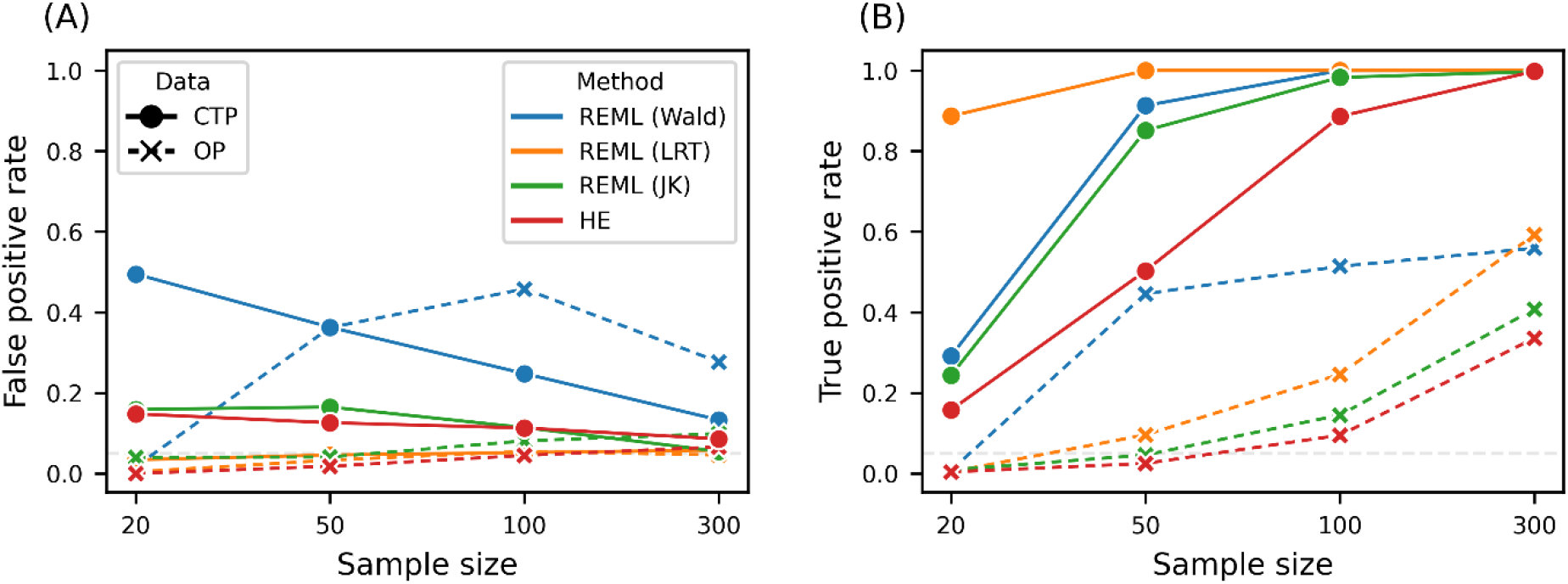
Power of CTMM’s test of cell type-specific variance in simulations with varying sample sizes. (A) In the simulation of Hom model, there is no cell type-specific variance; (B) in the simulation of Free model, each cell type has its own cell type-specific variance.

To examine the impact of uncertain estimates of *v_ic_*, we repeated the CTP simulation while incorporating noisy *v_ic_*. To be more realistic, this simulation was conducted with parameters estimated from real data of iPSCs differentiation. We first evaluated the uncertainty of *v_ic_* by bootstrap resampling cells for all combinations of individual, cell type, and gene. Most of them had a coefficient of variation around 0.2 (Figure S7). To incorporate this uncertainty into simulation, we added noise into *v_ic_* when fitting models (Methods). We tried five distributions of noise to cover the distribution of coefficient of variation in real data (Figure S7). In the simulation of Hom model, when there was no noise of *v_ic_*, that is fitting model with real *v_ic_* used in simulations, REML (LRT) and REML (JK) were well calibrated, HE was slightly inflated (Figure 2A). Along with the increase of noise, REML (LRT)’s false positive rate increased quickly and reached ~80% when using a high level of noise (coefficient of variation = 0.45); REML (JK) was rather resistant to noise that it only completely broke when unrealistically strong noise was added; while HE was not impacted by noise, it remained slightly inflated for all levels of noise. Of note, estimates of cell type-specific variance were weakly biased in REML under strong noise (Supplementary Figure S8). In the simulation of Free model, we found that REML (JK) had 80% of positive rate even when the cell type-specific variance is weak with 0.05 variance for the first cell type; HE also had good power with about 50% of positive rate when the first cell type had 0.05 variance (Figure 2B). Taken together, REML (JK) is the most powerful and robust method and is our primary approach in our iPSCs analysis.

**Figure 2.**
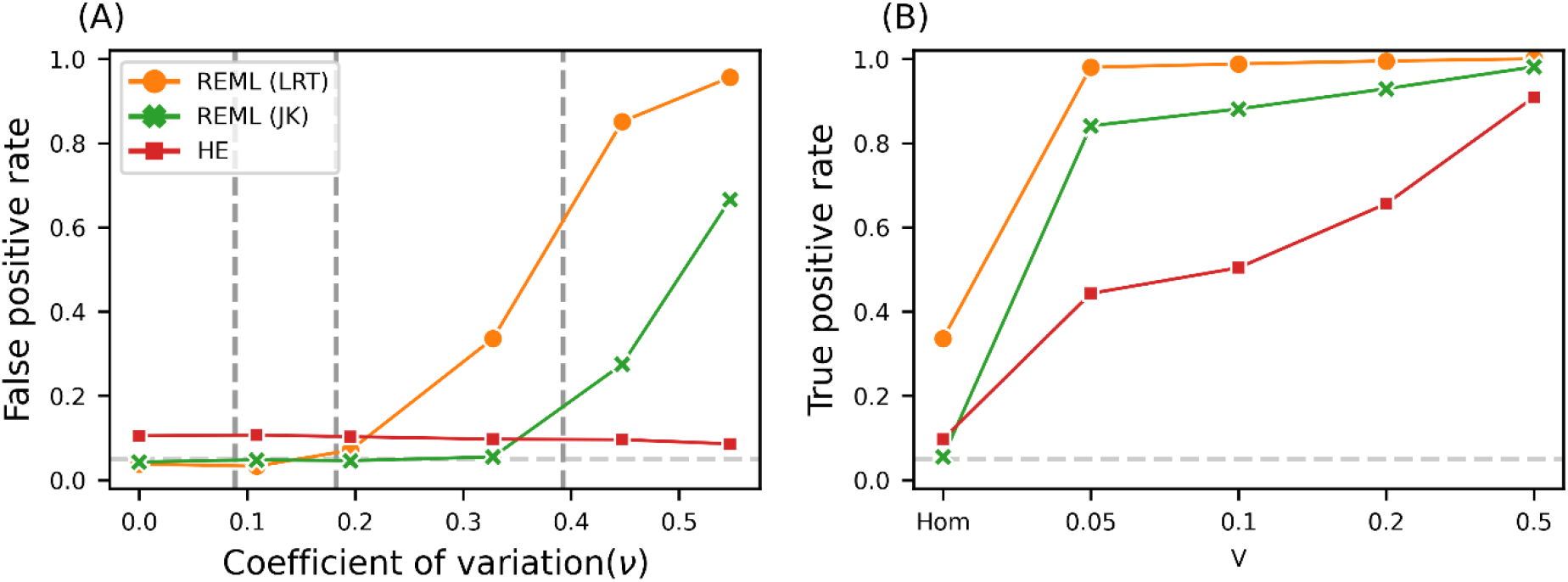
Power of CTMM in simulations with uncertain estimates of noise variance (*v_ic_*). (A) false positive rate under different levels of noise of *v_ic_*. Dashed lines indicating the 10%, 50%, and 90% percentiles of the transcriptome-wide distribution of coefficient of variation for *v_ic_* in the real iPSCs data; (B) true positive rate under noise of *v_ic_* with a coefficient of variation of 0.33, with varying cell type-specific variance for the first cell type from 0.05 to 0.5; cell type-specific variance for other three cell types were fixed to 0.1; in Hom, all cell types had 0 cell type-specific variance.

### Application to human induced pluripotent stem cells

We applied our methods to differentiating iPSCs^15^. Before fitting CTMM, we compared different approaches to imputing the cell type-specific pseudobulk (*y_ic_*, that is CTP). We evaluated single-gene imputation with softImpute and MVN and transcriptome-wide imputation with softImpute. We found that transcriptome-wide softImpute performed best (Figure S9 A, C), though MVN performed similarly. We also compared approaches to impute the noise variance (*v_ic_*), which is required for CTMM. We observed similar results as for the pseudobulk in terms of mean squared error (Figure S9 B, D). Based on these results, we used transcriptome-wide softImpute in practice.

We fit the Free model with both OP and CTP data using ML, REML, and HE (Figures S10 and S11). We focus on REML with CTP, which was most powerful and robust in simulations. Transcriptome-wide, we found that the variation across individuals was almost entirely cell type-specific, as the homogeneous variance has median close to 0 (*median* = 0.4%, Figure 3A). By contrast, cell type-specific variance has median 32.8% across cell types. Accounting for cell type proportions, cell type-specific variation explained 14% of interindividual expression variation transcriptome-wide (Supplementary Note Eq 3). Additionally, cell type-specific mean differences explained 12.3%, and measurement noise explained 9.5%. This illustrates the importance of modeling cell type-specific expression.

**Figure 3.**
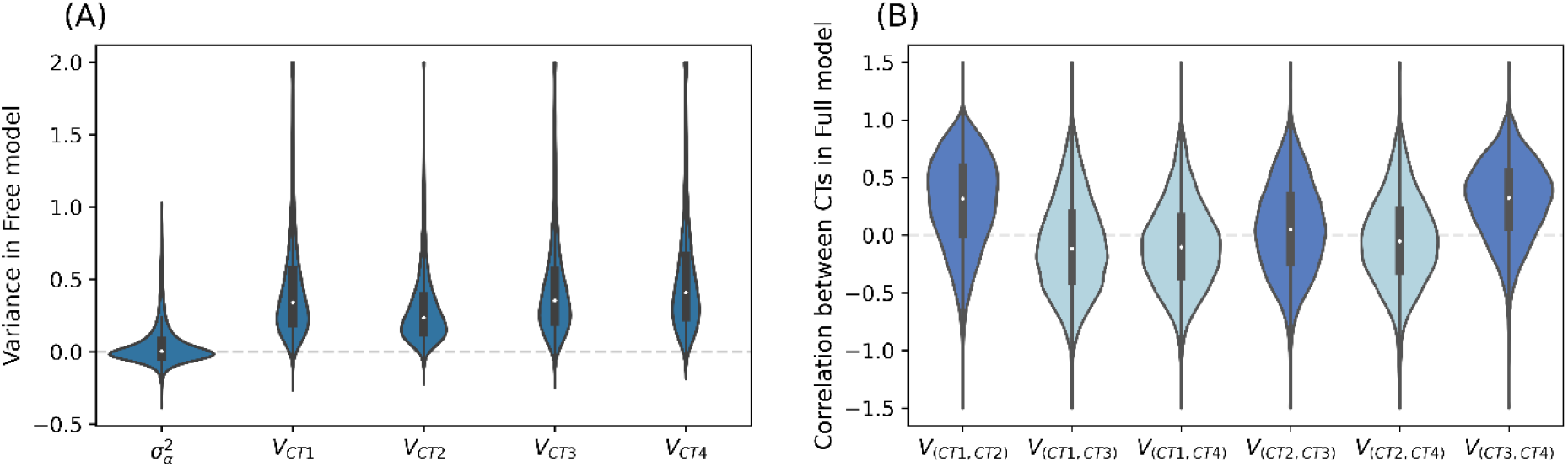
Distribution of variance and correlation of cell type-specific effect across transcriptome from REML with CTP data. (A) homogeneous variance 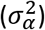 shared across cell types and cell type-specific variance from Free model; (B) correlation of cell type-specific effect between cell types from Full model, with dark blue indicating pairs of adjacent cell types and light blue indicating others. In (A), values above 2 were truncated; in (B), values above 1.5 or below −1.5 were truncated. 109 genes were excluded from (B) because of negative cell type-specific variance.

To evaluate the correlation of gene expression between cell types, we next fit the Full model. As expected, the correlations between adjacent development stages (CT1 and CT2, CT2 and CT3, and CT3 and CT4) were larger than the correlations between more distant stages (Figure 3B). Furthermore, the correlation between CT2 and CT3 (median: 0.050) was smaller than the other adjacent stages (CT1-CT2 median: 0.317; CT3-CT4 median: 0.323). This is consistent with rapid changes in molecular profiles at day2 (CT3)^15^. These patterns were also observed when fitting with ML or HE (Supplementary Figure S10). When fitting OP, as expected, the estimates were far less precise, especially for HE (Supplementary Figure S11).

Figure 4 shows gene expression differentiation in mean and variance in fitting CTP with REML (JK). We found many genes that were differentiated in variance between cell types, that is at least one cell type with non-zero cell type-specific variance. Among them, the top gene *POU5F1* (Wald *p* = 1.77 × 10^-27^), also known as OCT4, is one of the three core transcription factors in the pluripotency gene regulatory network^25^. This signal was alco confirmed in HE with CTP, where *POU5F1* was the most significant signal in variance differentiation (*p* = 1.67 × 10^-28^, Supplementary Figure S12). Although this gene was also significantly differentiated in mean, it is not outstanding in either REML or HE and less likely to be discovered for further functional analyses. To control for false positive, we identified candidate genes for cell type-specific variance as ordered by *p* value in REML (JK) meanwhile requiring significant signals after Bonferroni correction in both REML (JK) and HE, with top 10 genes shown in Table 1. Among them, 61 genes were not differentiated in mean (*p* > 0.01 in REML), with some of those genes involved in processes like cell differentiation and growth (see top 10 of those genes in Table 2). Take *NDUFB4* for example, there was no differentiation in mean between cell types (*p* = 0.014 in REML and *p* = 0.015 in HE), while significant differentiation in variance in both REML (*p* = 1.7 × 10^-19^) and HE (*p* = 8.59 × 10^-15^) (Figure 5). Of note, there were three marker genes used in *Cuomo et al.*^15^ to indicate each stem cell differentiation stage, spanning iPSC (*NANOG*), mesendoderm (*T*), and definitive endoderm (*GATA6*). We successfully detected significant mean differentiation in all three marker genes in both REML and HE; on the other hand, we detected significant variance differentiation in all three genes in HE, while only in *NANOG* gene in REML, indicating loss of power in REML. We also note that mean differentiation had much stronger signals than variance differentiation in both REML and HE (Supplementary Figure S13). We compared *p* values for variance differentiation from different tests when fitting CTP (Supplementary Figure S14). Generally, *p* values from different tests were largely consistent, except for REML with LRT. Specifically, for REML (JK) and HE, there were 4,773 genes that were significant in both; 341 genes were significant only in HE, likely to be false positive; 2,003 genes were significant only in REML (JK), partially due to false positive and partially due to higher power in REML (JK) than HE. We also conducted tests with OP data. Consistent with low power observed in simulations, we identified 23 genes that were significantly differentiated in variance in REML (LRT), and 0 genes were identified in HE (Supplementary Figure S15).

**Figure 4.**
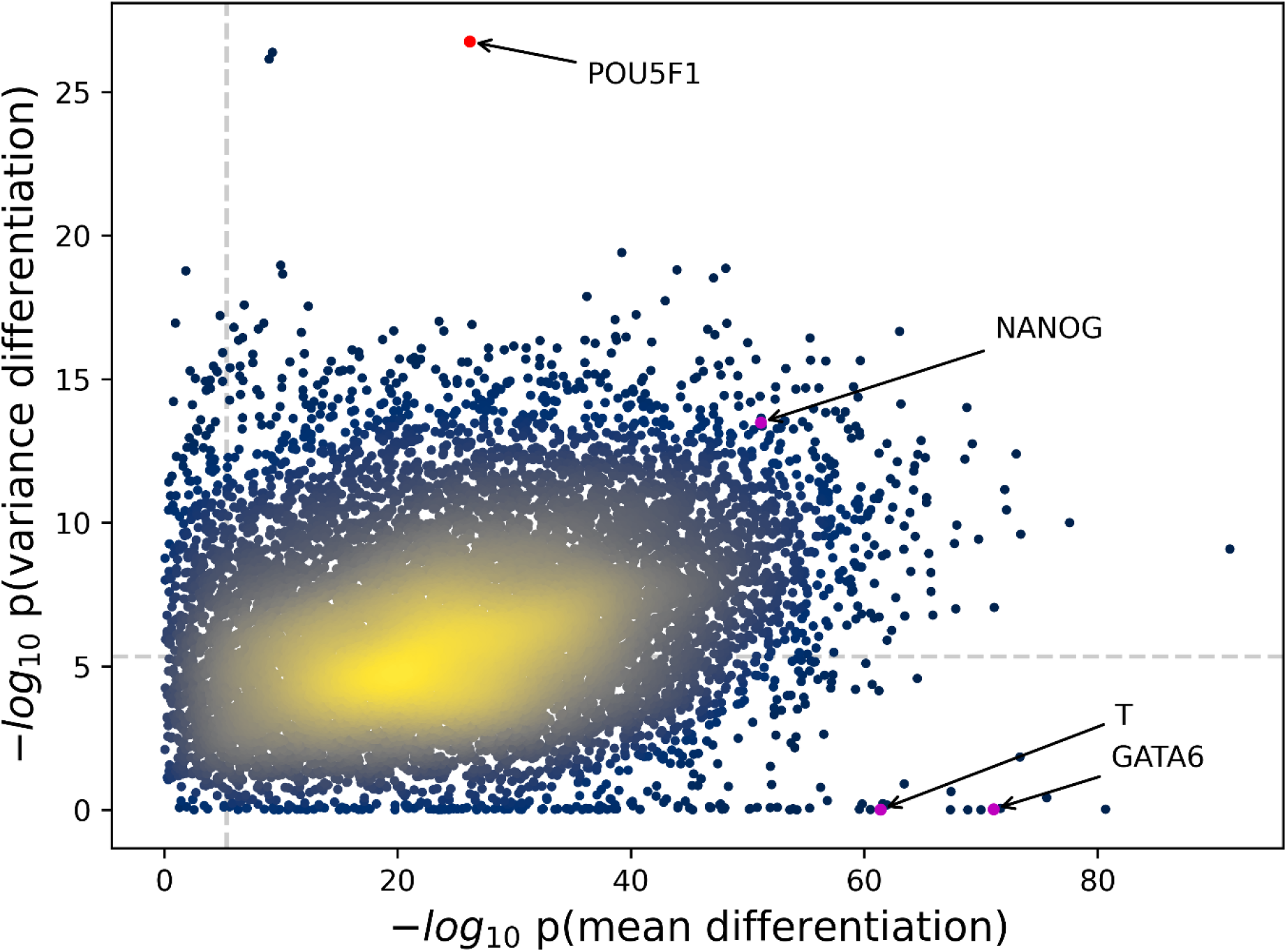
Distribution of *p* values for differentiation in expression mean and variance across transcriptome using REML with jackknife with CTP data. Each dot represents a gene. Dots are colored by the density of genes in the area, with yellow indicating denser distribution. Dashed lines indicate significance threshold after Bonferroni correction. The three purple dots indicate the three maker genes used in *Cuomo et al.* to indicate each stem cell differentiation stage, spanning iPSC (*NANOG*), mesendoderm (*T*), and definitive endoderm (*GATA6*). The red dot indicates the top signal *POU5F1,* which is one of the three core regulators in cell pluripotency.

**Figure 5.**
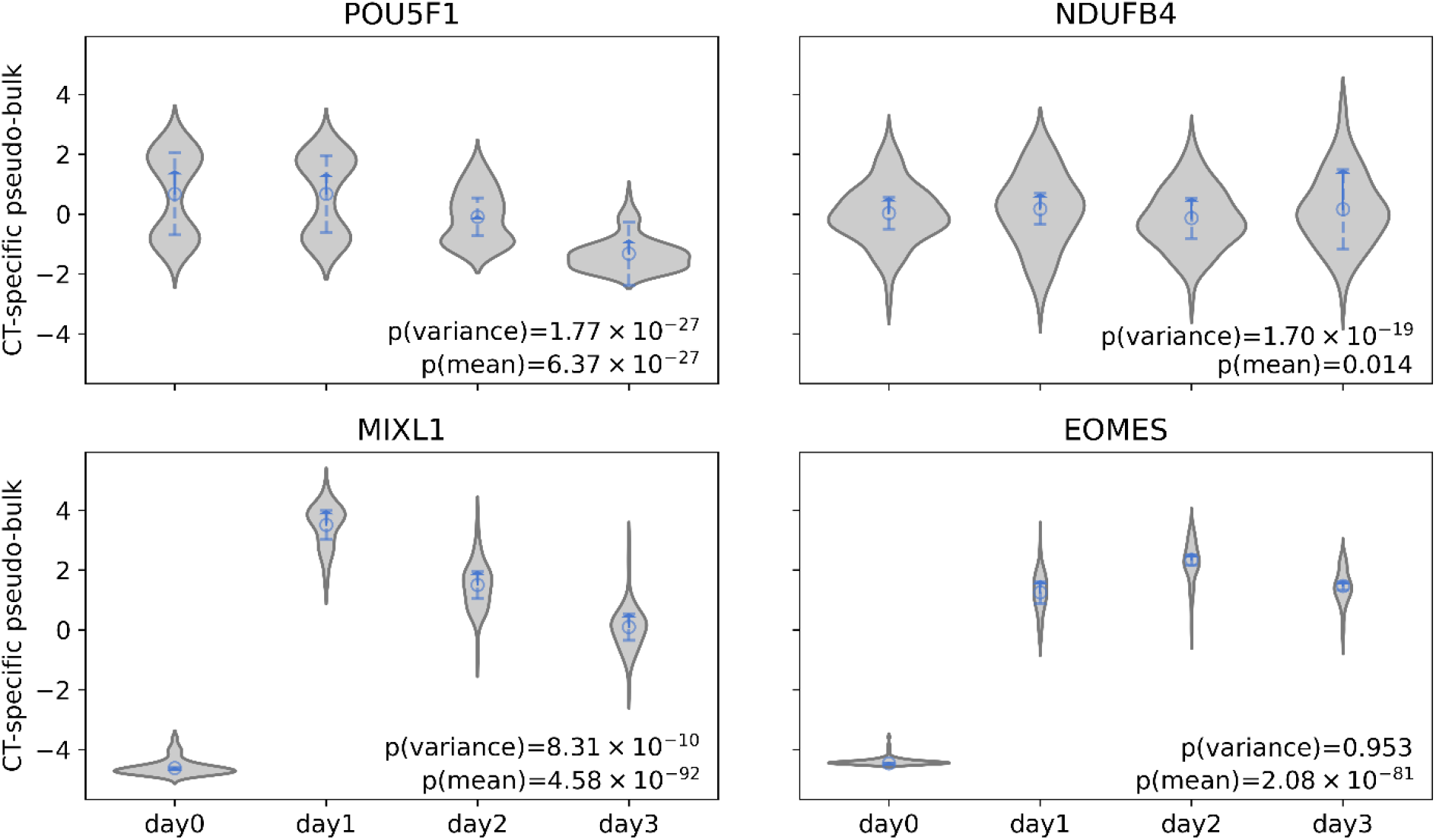
Estimates of cell type fixed effect and cell type-specific variance for specific genes in REML with CTP data. Four genes were chosen as examples. *POU5F1* exhibited the strongest signal of variance differentiation among all genes; *NDUFB4* exhibited the strongest signal of variance differentiation among genes without signals of mean differentiation (*p* > 0.01); *MIXL1* exhibited the strongest signal of mean differentiation among all genes; and *EOMES* exhibited the strongest signal of mean differentiation among genes without signals of variance differentiation (*p* > 0.01). The violin plot represents the distribution of cell type-specific pseudobulk after standardizing overall pseudobulk to mean 0 and variance 1; circles indicate estimated cell type fixed effects; the length of dash line indicates estimated cell type-specific variance; the length of arrow indicates the sum of homogeneous variance shared across cell types and cell type-specific variance.

**Table1.** Top 10 genes significantly differentiated in variance between cell types in REML, while remaining significant in HE, with CTP data.

| Gene | REML |  | HE |  | Function |
| --- | --- | --- | --- | --- | --- |
| | $p(\text{variance})^a$ | $p(\text{mean})^b$ | $p(\text{variance})$ | $p(\text{mean})$ | |
| POU5F1 | $1.77 \times 10^{-27}$ | $6.37 \times 10^{-27}$ | $1.67 \times 10^{-28}$ | $1.75 \times 10^{-27}$ | Cell pluripotency<sup>25</sup> |
| HSPA8 | $3.92 \times 10^{-20}$ | $6.07 \times 10^{-40}$ | $1.37 \times 10^{-19}$ | $7.25 \times 10^{-40}$ | Cell pluripotency<sup>26</sup> |
| SUB1 | $1.09 \times 10^{-19}$ | $1.03 \times 10^{-10}$ | $5.76 \times 10^{-16}$ | $1.45 \times 10^{-10}$ | Cell differentiation<sup>27</sup> |
| NOP16 | $1.39 \times 10^{-19}$ | $7.41 \times 10^{-49}$ | $8.67 \times 10^{-15}$ | $4.54 \times 10^{-49}$ | Cell growth<sup>28</sup> |
| NDUFB4 | $1.7 \times 10^{-19}$ | 0.014 | $8.59 \times 10^{-15}$ | 0.015 | Cell differentiation<sup>29</sup> |
| RPL35 | $2.21 \times 10^{-19}$ | $7.08 \times 10^{-11}$ | $2.36 \times 10^{-20}$ | $7.18 \times 10^{-11}$ | Cell proliferation and survival<sup>30</sup> |
| CCND1 | $3 \times 10^{-19}$ | $8.27 \times 10^{-48}$ | $3.55 \times 10^{-15}$ | $7.07 \times 10^{-48}$ | Cell differentiation<sup>31</sup> |
| GYPC | $1.33 \times 10^{-18}$ | $5.92 \times 10^{-37}$ | $1.08 \times 10^{-16}$ | $1.45 \times 10^{-36}$ | Cell differentiation<sup>32</sup> |
| PTMA | $1.88 \times 10^{-18}$ | $1.14 \times 10^{-43}$ | $1.52 \times 10^{-16}$ | $2.17 \times 10^{-43}$ | Apoptosis<sup>33</sup> |
| SHFM1 | $2.64 \times 10^{-18}$ | $1.37 \times 10^{-7}$ | $1.15 \times 10^{-16}$ | $1.11 \times 10^{-7}$ | Cell pluripotency<sup>34</sup> |

<sup>a</sup> $p(\text{variance})$ indicates $p$ values for variance differentiation between cell types; <sup>b</sup> $p(\text{mean})$ indicates $p$ values for mean differentiation between cell types.

**Table 2.** Top 10 genes significantly differentiated between cell types in expression variance while not in mean (*p* > 0.01) in REML, meanwhile significantly differentiated in variance in HE, with CTP data.

| Gene | REML |  | HE |  | Function |
| --- | --- | --- | --- | --- | --- |
| | $p(\text{variance})^a$ | $p(\text{mean})^b$ | $p(\text{variance})$ | $p(\text{mean})$ | |
| NDUFB4 | $1.7 \times 10^{-19}$ | 0.014 | $8.59 \times 10^{-15}$ | 0.015 | Cell differentiation<sup>29</sup> |
| NUTF2 | $1.12 \times 10^{-17}$ | 0.109 | $1.29 \times 10^{-16}$ | 0.105 | Cell cycle<sup>35</sup>, apoptosis<sup>36</sup> |
| SLX1A | $6.06 \times 10^{-15}$ | 0.162 | $7.11 \times 10^{-18}$ | 0.152 | Genome stability<sup>37</sup> |
| EIF4A1 | $2.4 \times 10^{-13}$ | 0.011 | $2.17 \times 10^{-10}$ | 0.044 | Stem cell self-renewal<sup>38</sup> |
| ATP5J2 | $4.96 \times 10^{-13}$ | 0.092 | $1.13 \times 10^{-13}$ | 0.084 | ATP synthesis<sup>39</sup> |
| TFPI2 | $5.81 \times 10^{-13}$ | 0.025 | $9.53 \times 10^{-9}$ | 0.016 | Cell proliferation<sup>40</sup> |
| SMAP1 | $1.07 \times 10^{-12}$ | 0.025 | $5.9 \times 10^{-10}$ | 0.026 | Cell growth<sup>41</sup> |
| NDN | $1.83 \times 10^{-12}$ | 0.011 | $7.75 \times 10^{-10}$ | 0.012 | Cell growth<sup>41</sup> |
| KRT10 | $2.38 \times 10^{-12}$ | 0.217 | $1.18 \times 10^{-10}$ | 0.214 | |
| NDUFA1 | $3.82 \times 10^{-12}$ | 0.406 | $2.71 \times 10^{-11}$ | 0.395 | Cell differentiation<sup>29</sup> |

<sup>a</sup> $p(\text{variance})$ indicates $p$ values for variance differentiation between cell types; <sup>b</sup> $p(\text{mean})$ indicates $p$ values for mean differentiation between cell types.

## Discussion

Mean differences in gene expression across cell types are well documented and are the primary focus of most scRNA-seq analyses. Here, we have introduced a new model called CTMM to quantify variance differences across cell types in scRNA-seq data. Bulk expression analyses have established that interindividual variance in expression can be important for characterizing disease biology^42^ and identifying context-dependent genetic effects^43^. The key innovation in CTMM is adapting Gaussian linear mixed models (LMMs) to scRNA-seq data, which is challenging because scRNA-seq data are highly noisy and non-Gaussian. The key idea is to summarize the scRNA-seq data into cell type-specific pseudobulk^17^, which enables approximately unbiased inference with CTMM on as few as 20 individuals. We carefully profile several standard methods to fit LMMs and propose a jackknife-based test using REML as the most powerful and robust method, which we support with extensive simulations and analyses of differentiating iPSCs. We implement and freely release these methods as a user-friendly Python package. We expect that CTMM will be an important step toward robust and rich variance decompositions of scRNA-seq data, which will be increasingly powerful and informative as scRNA-seq sample sizes grow.

In the limiting case with infinite cells, when the measurement error is reduced to 0, CTMM simplifies to a typical LMM on bulk expression. In this case, CTMM with Overall Pseudobulk (OP) data is comparable to decomposing variance in bulk expression data using computationally-deconvolved cell type proportions. The significant benefit is that scRNA data provides much better estimates of cell type proportions, which can both reduce false positives and improve power. Likewise, CTMM with Cell Type-specific Pseudobulk (CTP) data becomes comparable to bulk analyses of sorted cells without the need for sorting pre-defined cell types. In practice, when the number of cells is limited, another significant benefit of CTMM over bulk analyses is the ability to distinguish biological variance across individuals from measurement error, which is especially important when measurement error varies across individuals, cell types, or experimental conditions. However, the disadvantage of CTMM compared to bulk is that it requires larger sample sizes, which is currently expensive.

CTMM has several important limitations. First, as scRNA data in individual cells is highly non-Gaussian, CTMM’s Gaussian assumption relies on combining many cells and the central limit theorem. In practice, we require >10 cells per individual-cell type pair, which limits CTMM to common cell types. A related concern is that lowly-expressed genes can be severely non-Gaussian, increasing the number of cells needed for the Gaussian approximation. Second, CTMM assumes cell types are already known. Our real data analysis solves this by defining cell types based on experimental days. However, most studies infer cell types directly from the scRNA-seq data, inducing some circularity; this is typically ignored^2,7,44^ yet will deflate estimates of cell type specific variance by construction. Third, CTMM assumes discrete cell types, whereas continuous cell types are more appropriate in some cases, e.g., when defined by pseudo-time^15,16,45^ or degree of IFN stimulation^17^. While incorporating continuous cell types is straightforward with overall pseudobulk data, it can only be expressed in cell type-specific pseudobulk data by discretizing the continuous cell types. Fourth, it is well-known that count data evince a complex mean-variance relationship, and studies have observed that the variance of gene expression across cells is dependent on mean expression^46^. Nonetheless, we find biologically plausible genes with significant differential variance but without significant differential mean, showing that modelling variance has utility beyond merely tagging mean signals. Finally, despite our use of careful nonparametric tests, our Free test for cell type-specificity remains slightly inflated, emphasizing the importance of biologically validating and replicating results.

CTMM opens the door to translating well-established LMM methodologies to scRNA-seq data. A key extension of CTMM is to quantify cell type-specific heritability of gene expression, which is typically more powerful than single-SNP tests of context-specific genetic regulation^17,47^. Because CTMM models cell-level noise, it can eliminate downward biases in heritability that are unavoidable in bulk expression data. Another extension is to jointly model covariance across both cell types and genes. For example, this enables identifying cell type-shared and -specific networks. This, too, is necessarily biased in bulk expression data, where covarying measurement errors will confound biologically meaningful networks. The Full model can also be extended to learn structured networks between cell types by leveraging penalized covariance estimates^48^ or by specifically tailoring it to a given application; for example, we could restrict *V* to be banded to capture temporal structure in the differentiating iPSCs. The long-term goal is to combine together these features into a comprehensive model of transcriptomic covariation across cells, cell types, individuals, and environments in order to understand genetic and nongenetic drivers of complex disease. Overall, we consider CTMM an important step on a long path to fully understanding the causes and consequences of variation within and between individuals in scRNA-seq data.

## Data availability

Processed single cell count data from iPSCs were downloaded from Zenodo: https://zenodo.org/record/3625024#.Xil-0y2cZ0s.

## Code availability

CTMM Python package and code used for all analyses in this paper is available at: https://github.com/Minhui-Chen/CTMM.

## Acknowledgements

This work was funded by K25HL157603. We thank the Center for Research Informatics and the Research Computing Center for providing the compute resources.

## Author contributions

M.C. developed statistical methodology, performed analyses, and wrote the manuscript. A.D. conceived and supervised the project and wrote the manuscript.

