## Supplementary Note for "A robust model for cell type-specific interindividual variation in single-cell RNA sequencing data"

#### 1 Modelling Overall Pseudobulk data

We start with the model for Overall Pseudobulk (OP) data, which sums expression over all measured cells for each individual. The model in Eq 2 in the main text can be written as

$$y = P\beta + \alpha + (I \bullet P)\mathrm{vec}\left(\Gamma^T\right) + \delta$$

Here, $y \in \mathbb{R}^{N\times 1}$ is the vector of overall pseudobulk expression, where $N$ is the number of donor individuals. $P \in \mathbb{R}^{N\times C}$ is the matrix of cell type proportions, where $C$ is the number of cell types. (Note that $C$ is assumed known and fixed, as is the assignment of cells to cell types.) $\beta \in \mathbb{R}^{C\times 1}$ is the vector of cell type fixed effects, which represent the mean expression level for each cell type. $\alpha \in R^{N\times 1}$ is the vector of average expression across individuals that is homogeneously shared across all cell types, which we treat as a random effect:

$$\alpha \sim \mathcal{N}\left(0, \sigma_\alpha^2 I\right)$$

where $I$ is the identity matrix.

$\Gamma \in \mathbb{R}^{N\times C}$ is the matrix of cell type-specific average expression across all individuals and cell types, which we also treat as a random effect:

$$\Gamma \sim \mathcal{MN}\left(0, I, V\right)$$

This is written as a matrix normal random variable, where $I$ is the row-covariance matrix and $V$ is the column-covariance matrix. (Equivalently, we assume that $\mathrm{vec}\left(\Gamma^T\right) \sim \mathcal{N}\left(0, I \otimes V\right)$, where $\mathrm{vec}\left(\right)$ concatenates columns of a matrix into a vector.) In words, this means that we assume $\Gamma$ is i.i.d. across individuals and has covariance $V$ across cell types.

$\bullet$ is the transposed Khatri-Rao product. Although it may seem complex, $(A \bullet B)$ is simply Kronecker product of corresponding rows of $A$ and $B$. This generalizes the standard interaction between two univariate features to two feature matrices, and is implicitly constructed in any linear regression involving interactions.

Finally, $\delta \in \mathbb{R}^{N\times 1}$ is the vector of measurement noise due to sampling a finite number of cells. We assume that $\delta \sim \mathcal{N}\left(0, D\right)$, where $D := \mathrm{diag}\left(\nu\right)$ and $\nu \in R^{N\times 1}$ is the vector of noise levels for overall pseudobulk noise. Equivalently, this can be written $\delta_i \overset{\text{ind}}{\sim} \mathcal{N}\left(0, \nu_i\right)$. In our derivations, we assume that $\nu$ is known. In practice, we calculate it by evaluating the variance across cells (see main text for details), but we explore violations of this assumption in simulations (see main text and Section 3.3 below).

In practice, it is often essential to include other covariates in the model as either fixed or random effects, such as experiment batch, age, and sex. For simplicity and statistical parsimony, we assume that the effects of these covariates are shared homogeneously across cell types. We can include these effects by slightly generalizing our above model:

$$y = Xb + \alpha + (I \bullet P)\text{vec}\left(\Gamma^T\right) + Z\mu + \delta$$

where $b$ is the vector of fixed effects with the design matrix $X$, which now includes both traditional covariates (e.g., age and sex) as well as the cell type fixed effects $\beta$, and $\mu$ is the vector of random effects with the design matrix $Z$, such as batch effects. It can be useful to partition $Z\mu$ as:

$$Z\mu = \begin{bmatrix} Z_1 & ... & Z_r \end{bmatrix} \begin{bmatrix} \mu_1 \\ ... \\ \mu_r \end{bmatrix} = \sum_{i=1}^{r} Z_i \mu_i$$

where $Z_i$ and $\mu_i$ are the design matrix and the random effect vector for feature $i$. We assume that all random effects are distributed $\mu_i \overset{\text{ind}}{\sim} \mathcal{N}\left(0, \sigma_i^2 I\right)$.

Let $\circ$ be the element-wise (or Hadamard) product, and let $\otimes$ be the Kronecker (or tensor) product. Under this more general model of OP data that includes random effects of a factor variable $Z$, the distribution of $y$ is:

$$\begin{aligned}
\mathbb{E}\left(y\right) =& Xb \\
\mathbb{V}\left(y\right) =& \mathbb{V}\left(\alpha\right) + \mathbb{V}\left((I \bullet P)\text{vec}\left(\Gamma^T\right)\right) + \sum \mathbb{V}\left(Z_i \mu_i\right) + \mathbb{V}\left(\delta\right) \\
=& \sigma_\alpha^2 I + (I \bullet P)\mathbb{V}\left(\text{vec}\left(\Gamma^T\right)\right)(I \bullet P)^T + \sum Z_i \mathbb{V}\left(\mu_i\right) Z_i^T + D \\
=& \sigma_\alpha^2 I + (I \bullet P)(I \otimes V)(I \bullet P)^T + \sum Z_i \sigma_i^2 Z_i^T + D \\
=& \sigma_\alpha^2 I + I \circ (PVP^T) + \sum_{i=1}^{r} \sigma_i^2 Z_i Z_i^T + D \\
\implies y \sim& \mathcal{N}\left(Xb, \sigma_\alpha^2 I + I \circ (PVP^T) + \sum_{i=1}^{r} \sigma_i^2 Z_i Z_i^T + D\right)
\end{aligned} \tag{1}$$

#### 1.1 Fitting Overall Pseudobulk with ML, REML, and HE

### ML

We fit maximum likelihood (ML) estimates of $\beta$, $\sigma_\alpha^2$, and $V$ by maximizing the log-likelihood function jointly over these parameters (as well as over the other fixed effects in $b$ and the random effect variance components $\sigma_i^2$):

$$l(b, \sigma_\alpha^2, V, \sigma_1^2, ..., \sigma_r^2 | y) = -\frac{1}{2}[const + \ln|\mathbb{V}\left(y\right)| + (y - Xb)^T (\mathbb{V}\left(y\right))^{-1}(y - Xb)]$$

where *const* is a constant and $\mathbb{V}\left(y\right)$ means the covariance matrix of the vector $y$ that depends on $\sigma_\alpha^2$ and $V$ (and $\sigma_i^2$), as shown in Eq 1.

#### REML

For restricted maximum likelihood (REML) estimates of $V$ (and $\sigma_i^2$), we instead maximize the restricted likelihood (Harville, 1974; Harville, 1977):

$$l(\sigma_\alpha^2, V, \sigma_1^2, ..., \sigma_r^2|y) = -\frac{1}{2}[const + \ln|\mathbb{V}(y)| + \ln|X^T\mathbb{V}(y)^{-1}X| + y^T Hy]$$

where $H$ is defined as:

$$H := \mathbb{V}(y)^{-1} - \mathbb{V}(y)^{-1}X(X^T\mathbb{V}(y)^{-1}X)^{-1}X^T\mathbb{V}(y)^{-1}$$

After fitting these variance components, we then fit the fixed effects in REML using generalized least squares.

For both ML and RMEL, we maximize the likelihood function using the BFGS algorithm implemented in the R function 'optim'. To initialize optimization, in ML, we initialize fixed effects with ordinary least square estimates and initialize variances of random effects with parameters such that each variance component explains an equal amount of residual OP variance after subtracting off the fixed effects; in REML, since fixed effect sizes are not involved in the likelihood function, we initialize variances of random effects with parameters such that each variance component explains an equal amount of OP variance. If the initial optimization attempt fails to converge or has one variance component that explains more than 5 folds of OP variance, we rerun optimization for 10 times with random initial parameters and pick the converged run with the largest likelihood as the final result, in order to mitigate bias from local maxima. We allow negative variance components to reduce bias, though the total expression variance is always positive.

## HE

For Haseman-Elston regression (HE), we estimate variance components by method-of-moments after residualizing out fixed effects. This residualization uses the projection matrix $M := I - X(X^TX)^{-1}X^T$:

$$y' := My = M[\alpha + (I \bullet P)\text{vec}\left(\Gamma^T\right) + Z\mu + \delta]$$

Note that $M$ is symmetric ($M^T = M$) and idempotent ($M^2 = M$).

The HE estimator of $\theta := (\sigma_\alpha^2, V_{11}, ..., V_{CC}, V_{12}, ..., V_{(C-1)(C)}, \sigma_1^2, ..., \sigma_r^2)$ is obtained by minimizing the squared error between the expected and observed sample covariance matrix, i.e., the standard method-of-moments:

$$\begin{aligned}
\widehat{\theta} =& argmin_\theta||y'y'^T - \mathbb{V}(y')||_F \\
=& argmin_\theta||Myy^TM^T - M\mathbb{V}(y)M||_F \\
=& argmin_\theta||Myy^TM - M[\sigma_\alpha^2 I + I \circ (PVP^T) + \sum_{i=1}^r \sigma_i^2 Z_i Z_i^T + D]M||_F \\
=& argmin_\theta||(Myy^TM - MDM) - M[\sigma_\alpha^2 I + I \circ (PVP^T) + \sum_{i=1}^r \sigma_i^2 Z_i Z_i^T]M||_F
\end{aligned}$$

To solve this, rewrite it as a linear function of $\theta$,

$$\begin{aligned}
&\text{vec}\left(M[\sigma_\alpha^2 I + I \circ (PVP^T) + \sum_{i=1}^{r} \sigma_i^2 Z_i Z_i^T]M\right) \\
&= \text{vec}\left(MIM\right)\sigma_\alpha^2 + \text{vec}\left(M[I \circ (PVP^T)]M\right) + \sum_{i=1}^{r} \text{vec}\left(MZ_iZ_i^TM\right)\sigma_i^2 \\
&= \text{vec}\left(M\right)\sigma_\alpha^2 + \text{vec}\left(M[I \circ (\sum_{m,n} V_{mn} P_{,m}(P_{,n})^T)]M\right) + \sum_{i=1}^{r} \text{vec}\left(MZ_iZ_i^TM\right)\sigma_i^2 \\
&= \text{vec}\left(M\right)\sigma_\alpha^2 + \text{vec}\left(\sum_{m,n} M[I \circ (P_{,m}(P_{,n})^T)]MV_{mn}\right) + \sum_{i=1}^{r} \text{vec}\left(MZ_iZ_i^TM\right)\sigma_i^2 \\
&= \text{vec}\left(M\right)\sigma_\alpha^2 + \sum_{m,n} \text{vec}\left(M[I \circ (P_{,m}(P_{,n})^T)]M\right)V_{mn} + \sum_{i=1}^{r} \text{vec}\left(MZ_iZ_i^TM\right)\sigma_i^2 \\
&= Q\theta
\end{aligned}$$

Now we use the standard OLS (ordinary least square) projection of $\text{vec}\left(Myy^TM - MDM\right)$ onto the span of $Q$ to estimate $\theta$, with $\widehat{\theta} = (Q^TQ)^{-1}Q^T\text{vec}\left(Myy^TM - MDM\right)$.

Here, $Q^T := \begin{bmatrix} \text{vec}\left(M\right)^T \\ \text{vec}\left(M[I \circ (P_{,1}(P_{,1})^T)]M\right)^T \\ \cdots \\ \text{vec}\left(M[I \circ (P_{,C}(P_{,C})^T)]M\right)^T \\ 2\text{vec}\left(M[I \circ (P_{,1}(P_{,2})^T)]M\right)^T \\ \cdots \\ 2\text{vec}\left(M[I \circ (P_{,C-1}(P_{,C})^T)]M\right)^T \\ \text{vec}\left(MZ_1Z_1^TM\right)^T \\ \cdots \\ \text{vec}\left(MZ_rZ_r^TM\right)^T \end{bmatrix}$. There is a factor of 2 because $V$ is symmetric.

In special cases, $Q$ can be simplified. In Hom model where $V = 0$, we have $\theta = (\sigma_\alpha^2, \sigma_1^2, ..., \sigma_r^2)$ and $Q^T = \begin{bmatrix} \text{vec}\left(M\right)^T \\ \text{vec}\left(MZ_1Z_1^TM\right)^T \\ \cdots \\ \text{vec}\left(MZ_rZ_r^TM\right)^T \end{bmatrix}$; in Free model where $V_{m,n} = 0$ when $m \neq n$, we have $\theta = (\sigma_\alpha^2, V_{11}, ..., V_{CC}, \sigma_1^2, ..., \sigma_r^2)$ and $Q^T = \begin{bmatrix} \text{vec}\left(M\right)^T \\ \text{vec}\left(M[I \circ (P_{,1}(P_{,1})^T)]M\right)^T \\ \cdots \\ \text{vec}\left(M[I \circ (P_{,C}(P_{,C})^T)]M\right)^T \\ \text{vec}\left(MZ_1Z_1^TM\right)^T \\ \cdots \\ \text{vec}\left(MZ_rZ_r^TM\right)^T \end{bmatrix}$

Note that this expression is far more efficient than naively solving the linear system of equations directly because it takes advantage of the implicit structure in the $Q$ matrix.

### 2 Modelling Cell Type-specific Pseudobulk data

The Cell Type-specific Pseudobulk (CTP) data uses the same model on single cells as the OP data. However, in CTP, each individual's scRNA data is collapsed into a vector where each entry sums over all cells in a given cell type, rather than a single number that sums over all cells in all cell types. Formally, the Eq 3 model in the main text can be written as:

$$Y = 1_N\beta^T + \alpha 1_C^T + \Gamma + \delta$$

As in the OP model, $N$ is the number of individuals, $C$ is the number of cell types, $\beta$ is the mean expression in each cell type (shared across individuals), $\alpha$ is the mean expression in each individual (shared across cell types), and $\Gamma$ are cell type-specific variations across individuals. $1_N$ is a vector of 1s with length $N$, and $1_C$ is a vector of 1s with length $C$. We assume the same random effect distributions on $\alpha$ and $\Gamma$ as in the OP data.

The CTP data is a matrix $Y \in R^{N\times C}$. The OP data is roughly equal to the average across columns of the CTP data, and this holds exactly in the special case where all cell types are uniformly sampled in all individuals.

$\delta$ is now slightly different than in the OP data, as it is an $N\times C$ matrix instead of an $N\times 1$ vector. The difference is analogous to the difference between $y$ in the OP data and $Y$: approximately, the OP $\delta$ can be thought of as averaging across the columns of the CTP data. Its distribution is given by $\delta_{ic} \overset{\text{ind}}{\sim} \mathcal{N}(0, \nu_{ic})$, where $\nu_{ic}$ is analogous to the $\nu_i$ above for OP data.

As in the OP above, we include extra covariates as either fixed or random effects, to represent other factors that are shared across cell types, such as experiment batch, age, and sex.

$$Y = A\zeta 1_C^T + 1_N\beta^T + \alpha 1_C^T + \Gamma + Z\mu 1_C^T + \delta$$

Here, $\zeta$ is the vector of fixed effects (except for cell type fixed effect), $A$ is the corresponding deign matrix. There is a vector $1_C^T$ because there are $C$ cell types per individual, and the effect is assumed to be shared across cell types. The same as in the OP, $\mu$ is the vector of random effects, except for overall random effect and cell type-specific random effect, $Z$ is the design matrix for random effects.

We define $y := \text{vec}\left(Y^T\right)$ for this section, but note that this is not the same as the OP data in $y$ in Section 1.

$$\begin{aligned}
y =& \text{vec}\left(1_C\zeta^T A^T\right) + \text{vec}\left(\beta 1_N^T\right) + \text{vec}\left(1_C\alpha^T\right) + \text{vec}\left(\Gamma^T\right) + \text{vec}\left(1_C\mu^T Z^T\right) + \text{vec}\left(\delta^T\right) \\
=&(A\otimes 1_C)\zeta + (1_N\otimes I_C)\beta + (I_N\otimes 1_C)\alpha + \text{vec}\left(\Gamma^T\right) + (Z\otimes 1_C)\mu + \text{vec}\left(\delta^T\right) \\
=&Xb + (I_N\otimes 1_C)\alpha + \text{vec}\left(\Gamma^T\right) + \sum_{i=1}^{r}(Z_i\otimes 1_C)\mu_i + \text{vec}\left(\delta^T\right)
\end{aligned}$$

with $X := \begin{bmatrix} A\otimes 1_C & 1_N\otimes I_C \end{bmatrix}$ and $b := \begin{bmatrix}\zeta \\ \beta\end{bmatrix}$.

The expectation and variance of $Y$ is:

$$
\begin{aligned}
\mathbb{E}(y) =& Xb \\
\mathbb{V}(y) =& \mathbb{V}\left((I_N \otimes 1_C)\alpha\right) + \mathbb{V}\left(\text{vec}\left(\Gamma^T\right)\right) + \sum_{i=1}^{r} \mathbb{V}\left((Z_i \otimes 1_C)\mu_i\right) + \mathbb{V}\left(\text{vec}\left(\delta^T\right)\right) \\
=& (I_N \otimes 1_C)(\sigma_\alpha^2 I_N)(I_N \otimes 1_C)^T + I_N \otimes V + \sum_{i=1}^{r}(Z_i \otimes 1_C)\sigma_i^2(Z_i \otimes 1_C)^T + D \\
=& (I_N I_N) \otimes (1_C 1_C^T)\sigma_\alpha^2 + I_N \otimes V + \sum_{i=1}^{r}(Z_i Z_i^T) \otimes (1_C 1_C^T)\sigma_i^2 + D \\
=& (I_N \otimes J_C)\sigma_\alpha^2 + I_N \otimes V + \sum_{i=1}^{r}[(Z_i Z_i^T) \otimes J_C]\sigma_i^2 + D \\
=& I_N \otimes (J_C \sigma_\alpha^2 + V) + \sum_{i=1}^{r}[(Z_i Z_i^T) \otimes J_C]\sigma_i^2 + D \\
& \implies \\
& y \sim \mathcal{N}\left(Xb, I_N \otimes (J_C \sigma_\alpha^2 + V) + \sum_{i=1}^{r}[(Z_i Z_i^T) \otimes J_C]\sigma_i^2 + D\right) \qquad (2)
\end{aligned}
$$

where $J_C$ is a $C \times C$ matrix of 1s.

#### 2.1 Fitting Cell Type-specific Pseudobulk with ML, REML, and HE

##### ML and REML

These approaches are identical to the OP section, except that the CTP likelihood (or restricted likelihood) from (2) is used in place of the OP likelihood from (1).

For both ML and REML, we maximize the likelihood function using the BFGS algorithm implemented in the R function 'optim'. To initialize optimization, in ML, we initialize fixed effects with ordinary least square estimates and initialize variances of random effects with parameters such that each variance component explains an equal amount of residual CTP variance after subtracting off the fixed effects; in REML, since fixed effect sizes are not involved in the likelihood function, we initialize variances of random effects with parameters such that each variance component explains an equal amount of CTP variance. If the initial optimization attempt failed to converges or has one variance component that explains more than 5 folds of OP variance, we rerun optimization for 10 times with random initial parameters and pick the converged run with the largest likelihood as the final result, in order to mitigate bias from local maxima. We allow negative variance components to reduce bias, though the total expression variance is always positive.

### HE

As for OP data, we first projected out fixed effect by $M := I - X(X^TX)^{-1}X^T$:

$$
y' := My = M[(I_N \otimes 1_C)\alpha + \text{vec}\left(\Gamma^T\right) + \sum_{i=1}^{r}(Z_i \otimes 1_C)\mu_i + \text{vec}\left(\delta^T\right)]
$$

The HE estimator of $\theta := (\sigma_\alpha^2, V_{11}, ..., V_{CC}, V_{12}, ..., V_{(C-1)(C)}, \sigma_1^2, ..., \sigma_r^2)$ is obtained by minimizing the squared error between the expected and observed sample covariance matrix:

$$\begin{aligned}
\widehat{\theta} =& argmin_\theta ||y'y'^T - \mathbb{V}(y')||_F \\
=& argmin_\theta ||Myy^TM^T - M\mathbb{V}(y)M||_F \\
=& argmin_\theta ||Myy^TM - M[I_N \otimes (J_C\sigma_\alpha^2 + V) + \sum_{i=1}^{r} \left((Z_iZ_i^T) \otimes J_C\right)\sigma_i^2 + D]M||_F \\
=& argmin_\theta ||(Myy^TM - MDM) - M[I_N \otimes (J_C\sigma_\alpha^2 + V) + \sum_{i=1}^{r} \left((Z_iZ_i^T) \otimes J_C\right)\sigma_i^2]M||_F
\end{aligned}$$

To solve this, rewrite it as a linear function of $\theta$. For convenience, we define $[i]$ as the row or column indexes in a matrix corresponding to individual $i$, that is from $(i-1) \times C + 1$ to $i \times C$; we define $L_{m,n}$ as a $C \times C$ matrix of zeros, except for the entry of $(m, n)$, which equals to one.

$$\begin{aligned}
&\text{vec}\left(M[I_N \otimes (J_C\sigma_\alpha^2 + V) + \sum_{i=1}^{r} \left((Z_iZ_i^T) \otimes J_C\right)\sigma_i^2]M\right) \\
&= \text{vec}\left(M(I_N \otimes J_C)M\right)\sigma_\alpha^2 + \text{vec}\left(M(I_N \otimes V)M\right) + \sum_{i=1}^{r} \text{vec}\left(M\left((Z_iZ_i^T) \otimes J_C\right)M\right)\sigma_i^2 \\
&= \text{vec}\left(M(I_N \otimes J_C)M\right)\sigma_\alpha^2 + \text{vec}\left(M(I_N \otimes \sum_{m,n}^{C} V_{m,n}L_{m,n})M\right) + \sum_{i=1}^{r} \text{vec}\left(M\left((Z_iZ_i^T) \otimes J_C\right)M\right)\sigma_i^2 \\
&= \text{vec}\left(M(I_N \otimes J_C)M\right)\sigma_\alpha^2 + \sum_{m,n}^{C} \text{vec}\left(M(I_N \otimes L_{m,n})M\right)V_{m,n} + \sum_{i=1}^{r} \text{vec}\left(M\left((Z_iZ_i^T) \otimes J_C\right)M\right)\sigma_i^2 \\
&= Q\theta
\end{aligned}$$

Now we use the standard OLS projection of $\text{vec}\left(Myy^TM - MDM\right)$ onto the span of $Q$ to estimate $\theta$, with $\widehat{\theta} = (Q^TQ)^{-1}Q^T\text{vec}\left(Myy^TM - MDM\right)$.

Here, $$Q^T := \begin{bmatrix} \text{vec}\left(M(I_N \otimes J_C)M\right)^T \\ \text{vec}\left(M\left(I_N \otimes L_{1,1}\right)M\right)^T \\ \cdots \\ \text{vec}\left(M\left(I_N \otimes L_{C,C}\right)M\right)^T \\ \text{vec}\left(M\left(I_N \otimes (L_{1,2} + L_{2,1})\right)M\right)^T \\ \cdots \\ \text{vec}\left(M\left(I_N \otimes (L_{C-1,C} + L_{C,C-1})\right)M\right)^T \\ \text{vec}\left(M\left((Z_1Z_1^T) \otimes J_C\right)M\right)^T \\ \cdots \\ \text{vec}\left(M\left((Z_rZ_r^T) \otimes J_C\right)M\right)^T \end{bmatrix}.$$

In special cases, $Q$ can be simplified. In Hom model where $V = 0$, we have $\theta = (\sigma_\alpha^2, \sigma_1^2, ..., \sigma_r^2)$

and $Q^T = \begin{bmatrix} \text{vec}\,(M(I_N \otimes J_C)M)^T \\ \text{vec}\left(M\left((Z_1Z_1^T)\otimes J_C\right)M\right)^T \\ \cdots \\ \text{vec}\left(M\left((Z_rZ_r^T)\otimes J_C\right)M\right)^T \end{bmatrix}$; in Free model where $V_{m,n} = 0$ when $m \neq n$, we have

$\theta = (\sigma_\alpha^2, V_{11}, ..., V_{CC}, \sigma_1^2, ..., \sigma_r^2)$ and $Q^T = \begin{bmatrix} \text{vec}\,(M(I_N \otimes J_C)M)^T \\ \text{vec}\,(M\,(I_N \otimes L_{1,1})\,M)^T \\ \cdots \\ \text{vec}\,(M\,(I_N \otimes L_{C,C})\,M)^T \\ \text{vec}\left(M\left((Z_1Z_1^T)\otimes J_C\right)M\right)^T \\ \cdots \\ \text{vec}\left(M\left((Z_rZ_r^T)\otimes J_C\right)M\right)^T \end{bmatrix}$.

#### 2.2 Efficient computation: No additional random effects

In this section, through linear algebra manipulation, we reduce computation complexity for ML and REML under the special case where there are no random effects (other than $\alpha$ and $\Gamma$).

In ML, the rate limiting steps are the calculation of $\ln|\mathbb{V}(y)|$ and $(y-Xb)^T\mathbb{V}(y)^{-1}(y-Xb)$; in REML, those are the calculation of $\ln|\mathbb{V}(y)|$, $X^T\mathbb{V}(y)X$, $y^T\mathbb{V}(y)^{-1}y$, and $X^T\mathbb{V}(y)^{-1}y$. Since the general form for $(y-Xb)^T\mathbb{V}(y)^{-1}(y-Xb)$, $X^T\mathbb{V}(y)X$, $y^T\mathbb{V}(y)^{-1}y$, and $X^T\mathbb{V}(y)^{-1}y$ is $B^T\mathbb{V}(y)^{-1}F$, where $B$ and $F$ are tall matrices with only one or a few columns, we only need to reduce the computation complexity of $\ln|\mathbb{V}(y)|$ and $B^T\mathbb{V}(y)^{-1}F$.

When there is no extra random effect factor, i.e., $r = 0$. Define $[i]$ as indices corresponding to individual $i$, and $D_{[i][i]} := \text{diag}(\nu_{i,})$. The variance matrix of $y$ and its inverse can be simplified

$$\begin{aligned} \mathbb{V}(y) =& I_N \otimes (J_C\sigma_\alpha^2 + V) + D \\ =& \oplus_{i=1}^N (J_C\sigma_\alpha^2 + V + D_{[i][i]}) \\ \mathbb{V}(y)^{-1} =& \oplus_{i=1}^N \left(J_C\sigma_\alpha^2 + V + D_{[i][i]}\right)^{-1} \end{aligned}$$

Here, the $\oplus$ is the diagnal matrix of blocks, that is $A \oplus B := \begin{bmatrix} A & 0 \\ 0 & B \end{bmatrix}$ and $\oplus_{i=1}^2 A := \begin{bmatrix} A & 0 \\ 0 & A \end{bmatrix}$. Therefore, we can efficiently calculate the terms in the likelihoods of ML and REML:

$$\begin{aligned} \ln|\mathbb{V}(y)| =& \ln|\oplus_{i=1}^N \left(J_C\sigma_\alpha^2 + V + D_{[i][i]}\right)| \\ =& \ln\prod_{i=1}^N |J_C\sigma_\alpha^2 + V + D_{[i][i]}| \\ =& \sum_{i=1}^N \ln|J_C\sigma_\alpha^2 + V + D_{[i][i]}| \end{aligned}$$

$$\begin{aligned} B^T\mathbb{V}(y)^{-1}F =& B^T\left(\oplus_{i=1}^N \left(J_C\sigma_\alpha^2 + V + D_{[i][i]}\right)^{-1}\right)F \\ =& \sum_{i=1}^N B_{[i]}^T \left(J_C\sigma_\alpha^2 + V + D_{[i][i]}\right)^{-1} F_{[i]} \end{aligned}$$

Therefore, the computation complexity reduces from $O(N^3C^3)$ to $O(NC^3)$ for ML and REML.

#### 2.3 Efficient computation: One additional random effects

When there is only one extra random effect factor, i.e., $r = 1$, and each individual belongs to only one level of the factor, i.e. for each row of $Z_1$, there is only one element of 1 and all others are 0. Assuming the matrix $Z_1$ has $K$ levels and is ordered by levels, and each level $k$ has $x_k$ individuals, so that

$$Z_1 = \oplus_{k=1}^{K} 1_{x_k} = \begin{bmatrix} 1_{x_1} & & ... & \\ & 1_{x_2} & ... & \\ ... & ... & ... & ... \\ & & ... & 1_{x_K} \end{bmatrix}$$

Here, $1_{x_k}$ is a vector of 1 of length $x_k$. Then, $Z_1 Z_1^T = \oplus_{k=1}^{K} J_{x_k}$, where $J_{x_k}$ is a matrix of 1 of shape $x_k \times x_k$. Define $\{k\}$ as row or column indexes corresponding to individuals in level $k$. The variance of $y$ and its inverse can be simplified

$$\begin{aligned} \mathbb{V}(y) =& I_N \otimes (J_C\sigma_\alpha^2 + V) + [(Z_1 Z_1^T) \otimes J_C]\sigma_1^2 + D \\ =& \oplus_{k=1}^{K} \left(I_{x_k} \otimes (J_C\sigma_\alpha^2 + V)\right) + [(\oplus_{k=1}^{K} J_{x_k}) \otimes J_C]\sigma_1^2 + D \\ =& \oplus_{k=1}^{K} \left(I_{x_k} \otimes (J_C\sigma_\alpha^2 + V)\right) + \oplus_{k=1}^{K} (J_{x_k} \otimes J_C\sigma_1^2) + D \\ =& \oplus_{k=1}^{K} \left(I_{x_k} \otimes (J_C\sigma_\alpha^2 + V) + J_{x_k \times C}\sigma_1^2 + D_{\{k\}\{k\}}\right) \\ \mathbb{V}(y)^{-1} =& \oplus_{k=1}^{K} \left(I_{x_k} \otimes (J_C\sigma_\alpha^2 + V) + J_{x_k \times C}\sigma_1^2 + D_{\{k\}\{k\}}\right)^{-1} \end{aligned}$$

Therefore, we can efficiently calculate the terms in the likelihoods of ML and REML:

$$\begin{aligned} \ln|\mathbb{V}(y)| =& \ln| \oplus_{k=1}^{K} \left(I_{x_k} \otimes (J_C\sigma_\alpha^2 + V) + J_{x_k \times C}\sigma_1^2 + D_{\{k\}\{k\}}\right)| \\ =& \ln \prod_{k=1}^{K} |\left(I_{x_k} \otimes (J_C\sigma_\alpha^2 + V) + J_{x_k \times C}\sigma_1^2 + D_{\{k\}\{k\}}\right)| \\ =& \sum_{k=1}^{K} \ln|\left(I_{x_k} \otimes (J_C\sigma_\alpha^2 + V) + J_{x_k \times C}\sigma_1^2 + D_{\{k\}\{k\}}\right)| \end{aligned}$$

$$\begin{aligned} B^T\mathbb{V}(y)^{-1} F =& B^T \left(\oplus_{k=1}^{K} \left(I_{x_k} \otimes (J_C\sigma_\alpha^2 + V) + J_{x_k \times C}\sigma_1^2 + D_{\{k\}\{k\}}\right)^{-1}\right) F \\ =& \sum_{k=1}^{K} B_{\{k\},}^T \left(I_{x_k} \otimes (J_C\sigma_\alpha^2 + V) + J_{x_k \times C}\sigma_1^2 + D_{\{k\}\{k\}}\right)^{-1} F_{\{k\},} \end{aligned}$$

These manipulations reduce the computational complexity from $O(N^3C^3)$ to $O(KN_b^3C^3)$ for ML and REML, where $N_b$ is the number of individuals per level.

### 3 Simulation

#### 3.1 Variance partition of overall pseudobulk

We parameterize our simulations in terms of interpretable variance components of the OP data, which we partition using Eq2 in the main text by:

$$\begin{aligned}
\mathbb{V}(y) &= \mathbb{V}(\mathbb{E}(y_i)) + \mathbb{E}(\mathbb{V}(y_i)) \\
&= \mathbb{V}(P_{i,}\beta) + \mathbb{E}\left(\sigma_\alpha^2 + P_{i,}VP_{i,}^T + \nu_i\right) \\
&= \beta^T S\beta + \sigma_\alpha^2 + \mathbb{E}\left(P_{i,}VP_{i,}^T\right) + \mathbb{E}(\nu_i) \\
&= \beta^T S\beta + \sigma_\alpha^2 + \mathbb{E}\left(\mathrm{tr}\left(P_{i,}VP_{i,}^T\right)\right) + \mathbb{E}(\nu_i) \\
&= \beta^T S\beta + \sigma_\alpha^2 + \mathrm{tr}\left(V\mathbb{E}\left(P_{i,}^T P_{i,}\right)\right) + \mathbb{E}(\nu_i) \\
&= \beta^T S\beta + \sigma_\alpha^2 + \mathrm{tr}\left(V\left(S + \pi\pi^T\right)\right) + \mathbb{E}(\nu_i) \\
&= \underbrace{\beta^T S\beta}_{\text{cell type-specific mean}} + \underbrace{\sigma_\alpha^2}_{\text{cell type-shared variation}} + \underbrace{\mathrm{tr}(VS) + \pi^T V\pi}_{\text{cell type-specific variation}} + \underbrace{\mathbb{E}(\nu_i)}_{\text{measurement noise}}
\end{aligned} \tag{3}$$

Here, $S := \frac{1}{N}P_d^T P_d$ is the covariance matrix for cell type proportions; $P_d := P - 1_N\pi^T$ is demeaned cell type proportion matrix; $\pi$ is the vector of mean cell type proportions. We partition the variance of OP into four components: cell type fixed effect, homogeneous random effect, cell type-specific random effect, and noise.

#### 3.2 Simulation of overall pseudobulk and cell type-specific pseudobulk

We set parameters for our simulation according to the four variance components of OP in (3). In the simulation of Free model, we assumed each of the four components explained 25% of variance, i.e., $\beta^T S\beta = \sigma_\alpha^2 = \mathrm{tr}(VS) + \pi^T V\pi = \mathbb{E}(\nu_i) = 0.25$. We simulated 100 individuals and 4 cell types. Cell type proportions for each individual ($P_{i,}$) were i.i.d. sampled from Dirichlet distribution $Dir(2,2,2,2)$, so each cell type has an expected proportion of 25%, i.e., $\pi^T = \begin{bmatrix}0.25 & 0.25 & 0.25 & 0.25\end{bmatrix}$. From Dirichlet distribution, we also calculate the covariance matrix ($S$) of cell type proportions. Assuming the ratio of fixed effects for the four cell types is $\beta_1 : \beta_2 : \beta_3 : \beta_4 = 8 : 4 : 2 : 1$, we calculated the fixed effects for each cell type $\beta$, with the constraint of $\beta^T S\beta = 0.25$. Assuming the ratio of cell type-specific variances is $V_{11} : V_{22} : V_{33} : V_{44} = 64 : 16 : 4 : 1$, we calculated the cell type-specific variance matrix $V$ with the constraint of $\mathrm{tr}(VS) + \pi^T V\pi = 0.25$. We sampled the noise variance for each individual from Gamma distribution $\nu_i \overset{\text{ind}}{\sim} \Gamma(k = \frac{25}{4}, \theta = 0.04)$, which has $\mathbb{E}(\nu_i) = k\theta = 0.25$ and $\mathbb{V}(\nu_i) = k\theta^2 = 0.01$. Assuming that the residual effect $\epsilon_{ics}$ for gene expression in $s$-th cell from cell type $c$ in individual $i$ (as described in Eq 1 in the main text) is i.i.d. across all cells for each individual, such that $\mathbb{V}(\epsilon_{ics}) = \sigma_i^2$. Based on this assumption, we calculated the noise variance for CTP in each individual-cell type pair as $\nu_{ic} = \frac{\nu_i}{P_{ic}}$. With these parameters, we generated OP and CTP using Eq 2 and Eq 3 in the main text.

The simulation process for the Hom and Full models is similar to the Free model, except for the variance decomposition of OP. In the Hom model, since there is no cell type-specific variation in the model, we assumed that homogeneously shared variation accounts for 50% of the variance of OP, i.e., $\sigma_\alpha^2 = 0.5$. In the Full model, since $V$ and $\sigma_\alpha^2$ are not jointly identified, we set $\sigma_\alpha^2 = 0$ and cell type-specific variation explained OP variance to 50%, i.e., $\operatorname{tr}(VS) + \pi^T V \pi = 0.5$. To account for the correlation of cell type-specific random effect between cell types, we not only set the ratio of cell type-specific variances of $V_{11} : V_{22} : V_{33} : V_{44} = 1 : 1 : 1 : 1$, but also set a correlation of 0.9 between nearby cell types and a correlation of 0.7 or 0.5 for others.

To test the performance of CTMM in various situations, we varied one parameter at a time, including sample size, cell type proportions, and cell type-specific variances. A list of tested parameters for each model is available in Supplementary Table S1. For the simulation of Hom and Free models, we fit the simulated OP and CTP data into the Hom and Free models with maximum likelihood (ML), restricted maximum likelihood (REML), and Haseman-Elston regression (HE). We then tested for cell type-specific variance with applicable Wald tests and likelihood-ratio tests (LRT), as described in the main text. For the simulation of Full model, we fit the simulated OP and CTP data into the Full model with ML, REML, and HE but did not perform hypothesis testing due to the statistical and computational complexity. We ran 1,000 replicates for each set of parameters.

#### 3.3 Simulation with noisy $\nu$

As $\nu_{ic}$ is unknown and is estimated from cell-to-cell variation in practice, we also performed simulations to assess CTMM's sensitivity to estimation errors in $\nu_{ic}$. We only examined CTP in this simulation, as it's far more powerful. As the simulation focuses on CTMM's utility in our real data analysis, we simulated Hom and Free models using parameters estimated in the iPSCs data. For each set of simulation parameters, we ran 1,000 replicates.

In the simulation of Hom model, we randomly drew a gene from iPSCs data and obtained its parameters ($\sigma_\alpha^2$ and $\beta$) estimated using REML with CTP data under the Hom model. If $\sigma_\alpha^2 < 0$, we set it to 0. Using cell type proportions $P$ and measurement noise variance $\nu_{ic}$ from the iPSCs data, we generated CTP using Eq 3 in the main text. To incorporate estimation error in $\nu_{ic}$, for each $\nu_{ic}$, we draw $x_{ic}$ i.i.d. from a $Beta(2, b)$ distribution and then add $+x_{ic}\nu_{ic}$ or $-x_{ic}\nu_{ic}$ before inputting $\nu_{ic}$ to CTMM. To cover the distribution of estimation error in iPSCs data, we simulated $b = 20, 10, 5, 3, 2$, to get corresponding coefficients of variation of 0.11, 0.20, 0.33, 0.45, and 0.55 for $\nu_i c$.

We simulated under the Free model with varying cell type-specific variances. We obtained the parameters $\sigma_\alpha^2$ and $\beta$ in the same way as in the simulation of Hom model. We varied the cell type-specific variance for cell type 1 ($V_{11}$) from 0.05 to 0.5 and fixed other cell type-specific variances to 0.1. Those values were chosen based on the distribution of estimated cell type-specific variances in the iPSC data in order to make the simulation more realistic. For simplicity, the Free model simulations always use $b = 5$ (the most realistic value) to add estimation error into $\nu_{ic}$.
