## Supplementary Figure for "A robust model for cell type-specific interindividual variation in single-cell RNA sequencing data"

**Table S1. Parameters used in simulations of the Hom, Free, and Full models.**

| Model | Sample size | Dirichlet distribution of cell type proportions | Proportions of variances explained by the four components | Ratio of cell type-specific random effect variance | Correlation of cell type-specific random effect |
|---|---|---|---|---|---|
| Hom (baseline) | 100 | (2, 2, 2, 2) | (0.25, 0.5, 0, 0.25) | NA | NA |
| Hom | **20** | (2, 2, 2, 2) | (0.25, 0.5, 0, 0.25) | NA | NA |
| Hom | **50** | (2, 2, 2, 2) | (0.25, 0.5, 0, 0.25) | NA | NA |
| Hom | **300** | (2, 2, 2, 2) | (0.25, 0.5, 0, 0.25) | NA | NA |
| Hom | 100 | **(0.5, 2, 2, 2)** | (0.25, 0.5, 0, 0.25) | NA | NA |
| Hom | 100 | **(1, 2, 2, 2)** | (0.25, 0.5, 0, 0.25) | NA | NA |
| Hom | 100 | **(4, 2, 2, 2)** | (0.25, 0.5, 0, 0.25) | NA | NA |
| Free (baseline) | 100 | (2, 2, 2, 2) | (0.25, 0.25, 0.25, 0.25) | 64:16:4:1 | NA |
| Free | **20** | (2, 2, 2, 2) | (0.25, 0.25, 0.25, 0.25) | 64:16:4:1 | NA |
| Free | **50** | (2, 2, 2, 2) | (0.25, 0.25, 0.25, 0.25) | 64:16:4:1 | NA |
| Free | **300** | (2, 2, 2, 2) | (0.25, 0.25, 0.25, 0.25) | 64:16:4:1 | NA |
| Free | 100 | **(0.5, 2, 2, 2)** | (0.25, 0.25, 0.25, 0.25) | 64:16:4:1 | NA |
| Free | 100 | **(1, 2, 2, 2)** | (0.25, 0.25, 0.25, 0.25) | 64:16:4:1 | NA |
| Free | 100 | **(4, 2, 2, 2)** | (0.25, 0.25, 0.25, 0.25) | 64:16:4:1 | NA |
| Free | 100 | (2, 2, 2, 2) | **(0.25, 0.10, 0.40, 0.25)** | 64:16:4:1 | NA |
| Free | 100 | (2, 2, 2, 2) | **(0.25, 0.20, 0.30, 0.25)** | 64:16:4:1 | NA |
| Free | 100 | (2, 2, 2, 2) | **(0.25, 0.30, 0.20, 0.25)** | 64:16:4:1 | NA |
| Free | 100 | (2, 2, 2, 2) | **(0.25, 0.40, 0.10, 0.25)** | 64:16:4:1 | NA |
| Full (baseline) | 100 | (2,2,2,2) | (0.25, 0.25, 0.25, 0.25) | 1:1:1:1 | CT1-CT2, CT2-CT3, CT3-CT4: 0.9<br>CT1-CT3, CT2-CT4: 0.7<br>CT1-CT4: 0.5 |
| Full | **20** | (2,2,2,2) | (0.25, 0.25, 0.25, 0.25) | 1:1:1:1 | CT1-CT2, CT2-CT3, CT3-CT4: 0.9<br>CT1-CT3, CT2-CT4: 0.7<br>CT1-CT4: 0.5 |
| Full | **50** | (2,2,2,2) | (0.25, 0.25, 0.25, 0.25) | 1:1:1:1 | CT1-CT2, CT2-CT3, CT3-CT4: 0.9<br>CT1-CT3, CT2-CT4: 0.7<br>CT1-CT4: 0.5 |
| Full | **300** | (2,2,2,2) | (0.25, 0.25, 0.25, 0.25) | 1:1:1:1 | CT1-CT2, CT2-CT3, CT3-CT4: 0.9<br>CT1-CT3, CT2-CT4: 0.7<br>CT1-CT4: 0.5 |

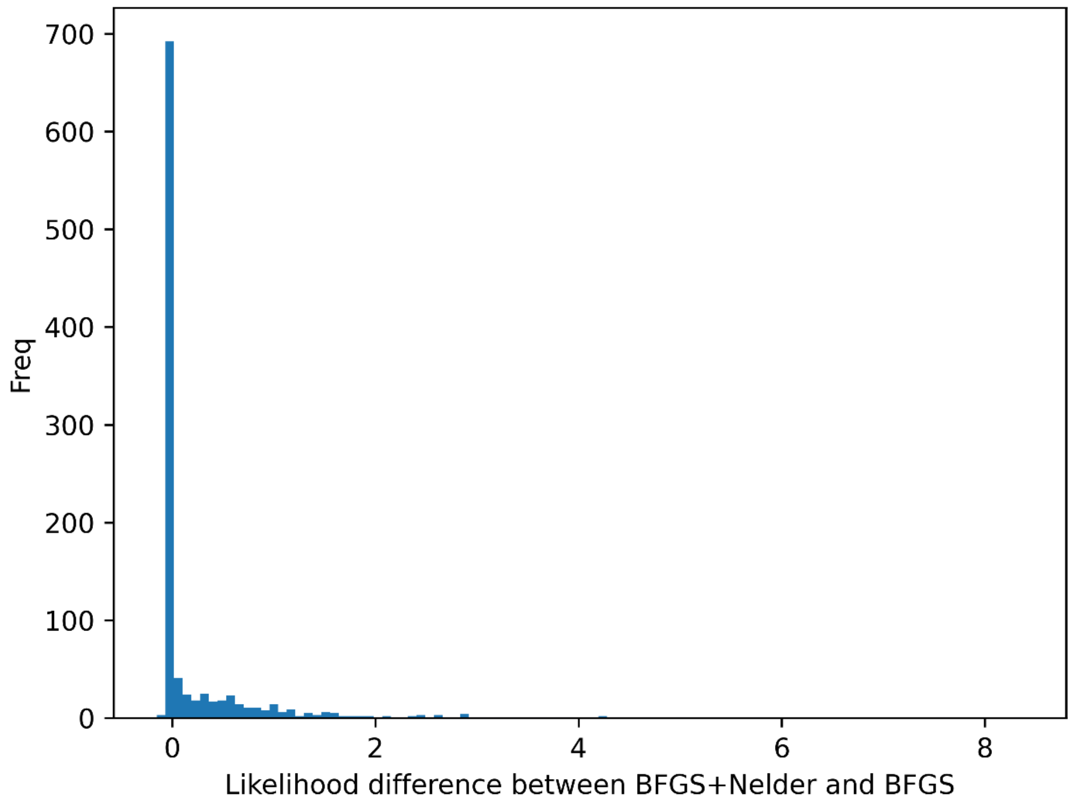


**Figure S1. Increase of inferred maximum likelihood in ML with OP after refining BFGS solution with Nelder-Mead algorithms.** The same simulated OP data from the Free baseline model (with parameters shown in Table S1) was fit by ML and optimized using BFGS alone or a combination of BFGS followed by Nelder-Mead. Both BFGS and Nelder-Mead algorithms are implemented using the R function 'optim'.

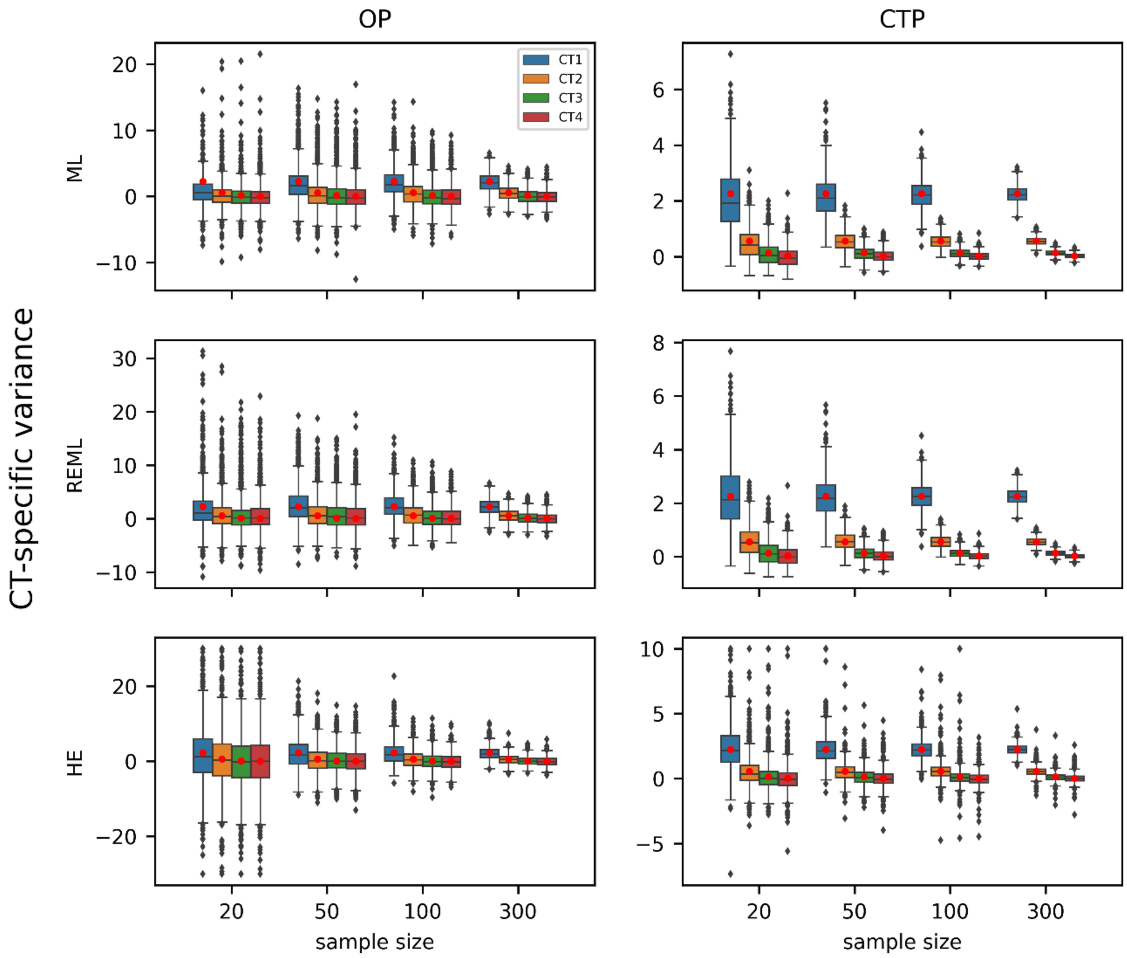


**Figure S2. CTMM estimates of cell type-specific variance with varying sample sizes.** Estimates are fit under the Free model, and red dots indicate the true cell type-specific variances. Rows show different estimation methods. Columns show different input to CTMM, either the overall pseudobulk (OP) or cell type-specific pseudobulk (CTP). Box plots indicate the distribution of estimated cell type-specific variances across 1,000 replicate simulations. In HE with OP, values above 30 or below -30 were truncated; in HE with CTP, values above 10 were truncated.

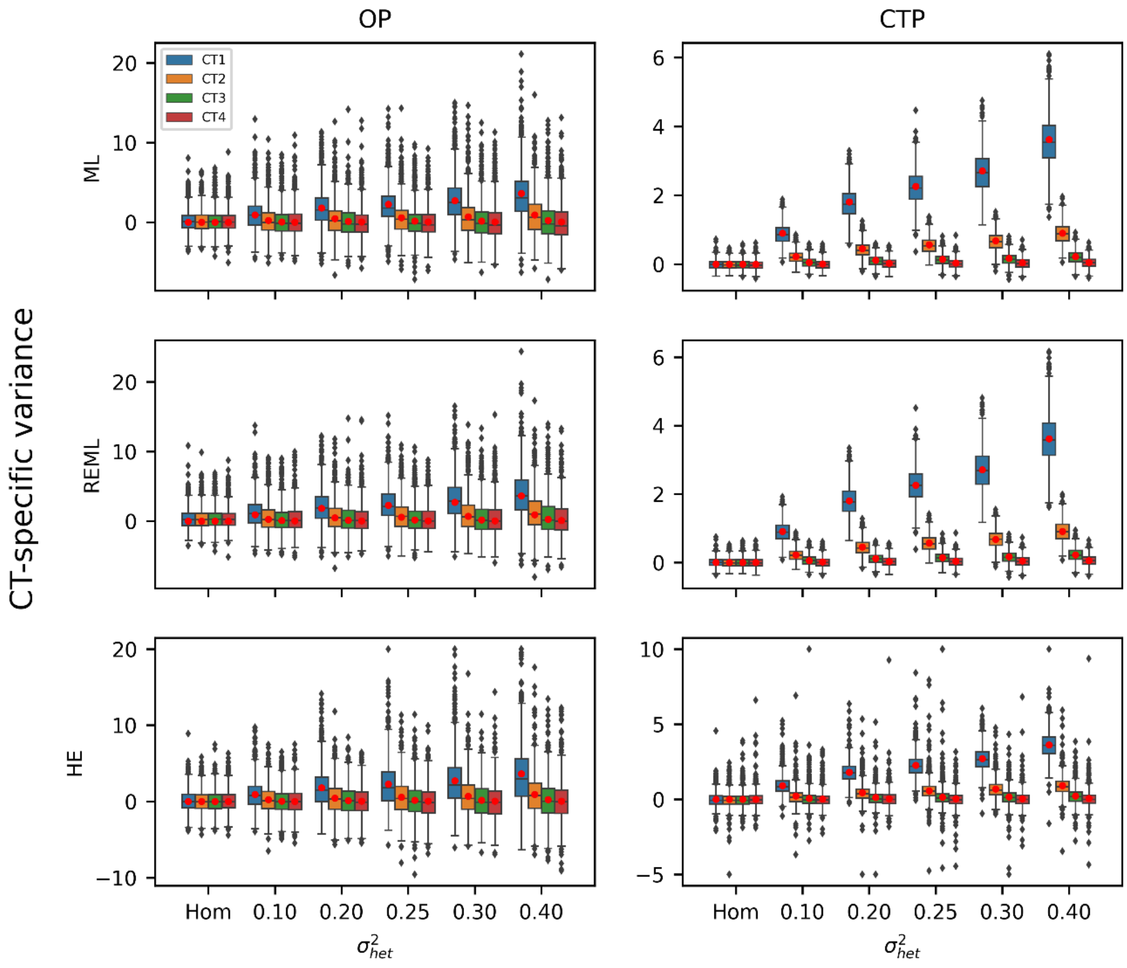


**Figure S3. CTMM estimates of cell type-specific variance with varying levels of true cell type-specific variance.** $\sigma^2_{het}$ represents the proportion of variance explained by cell type-specific variation when combined across all four cell types; the Hom model of no cell type-specificity is obtained when $\sigma^2_{het}$=0. Estimates are fit under the Free model, and red dots indicate the true cell type-specific variances. Rows show different estimation methods. Columns show different input to CTMM, either the overall pseudobulk (OP) or cell type-specific pseudobulk (CTP). Box plots indicate the distribution of estimated cell type-specific variances across 1,000 replicate simulations. In HE with OP, values above 20 were truncated; in HE with CTP, values above 10 or below -5 were truncated.

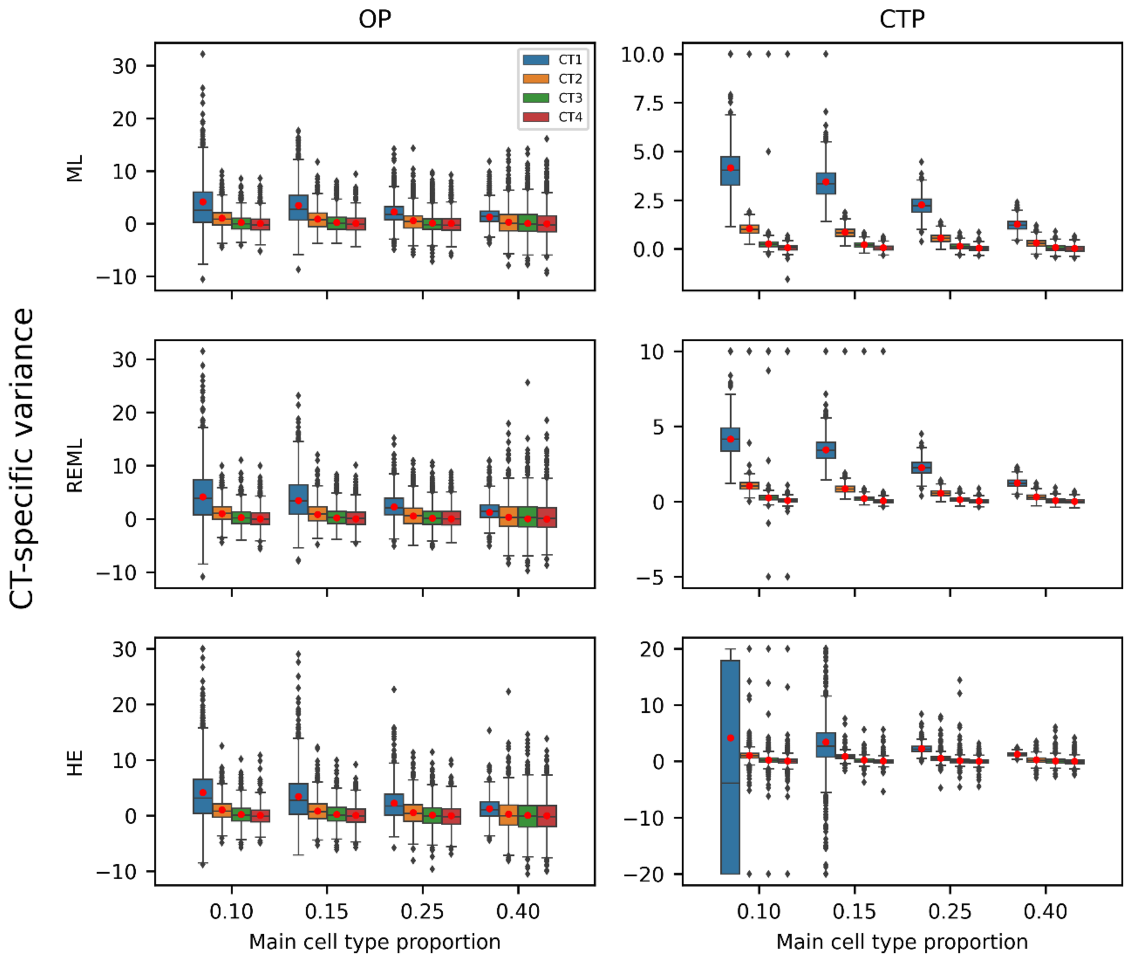


**Figure S4. CTMM estimates of cell type-specific variance with varying cell type proportions.** The proportion of the “main” cell type, which has the largest cell type-specific variance, is varied, with the proportions of other cell types scaled so the total proportions sum to 1. Estimates are fit under the Free model, and red dots indicate the true cell type-specific variances. Rows show different estimation methods. Columns show different input to CTMM, either the overall pseudobulk (OP) or cell type-specific pseudobulk (CTP). Box plots indicate the distribution of estimated cell type-specific variances across 1,000 replicate simulations. In HE with OP, values above 30 were truncated; in ML with CTP, values above 10 were truncated; in REML with CTP, values were truncated to (-5, 10); in HE with CTP, values were truncated to (-20, 20).

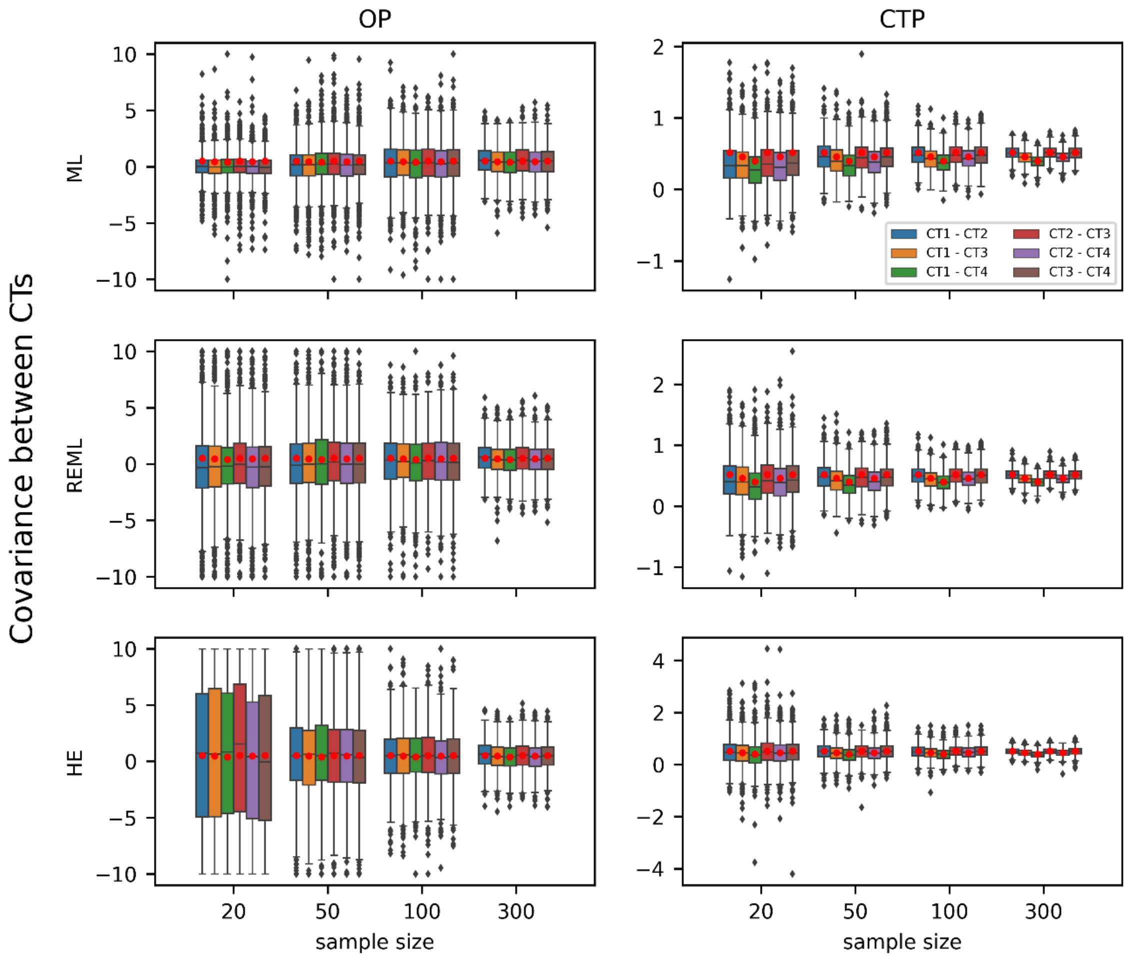


**Figure S5. CTMM estimates of cell type-specific covariance with varying sample sizes.** Estimates are fit under the Full model, and red dots indicate the true cell type-specific covariances. Rows show different estimation methods. Columns show different input to CTMM, either the overall pseudobulk (OP) or cell type-specific pseudobulk (CTP). Box plots indicate the distribution of estimated cell type-specific variances across 1,000 replicate simulations. in CTMM with OP, values were truncated to (-10, 10).

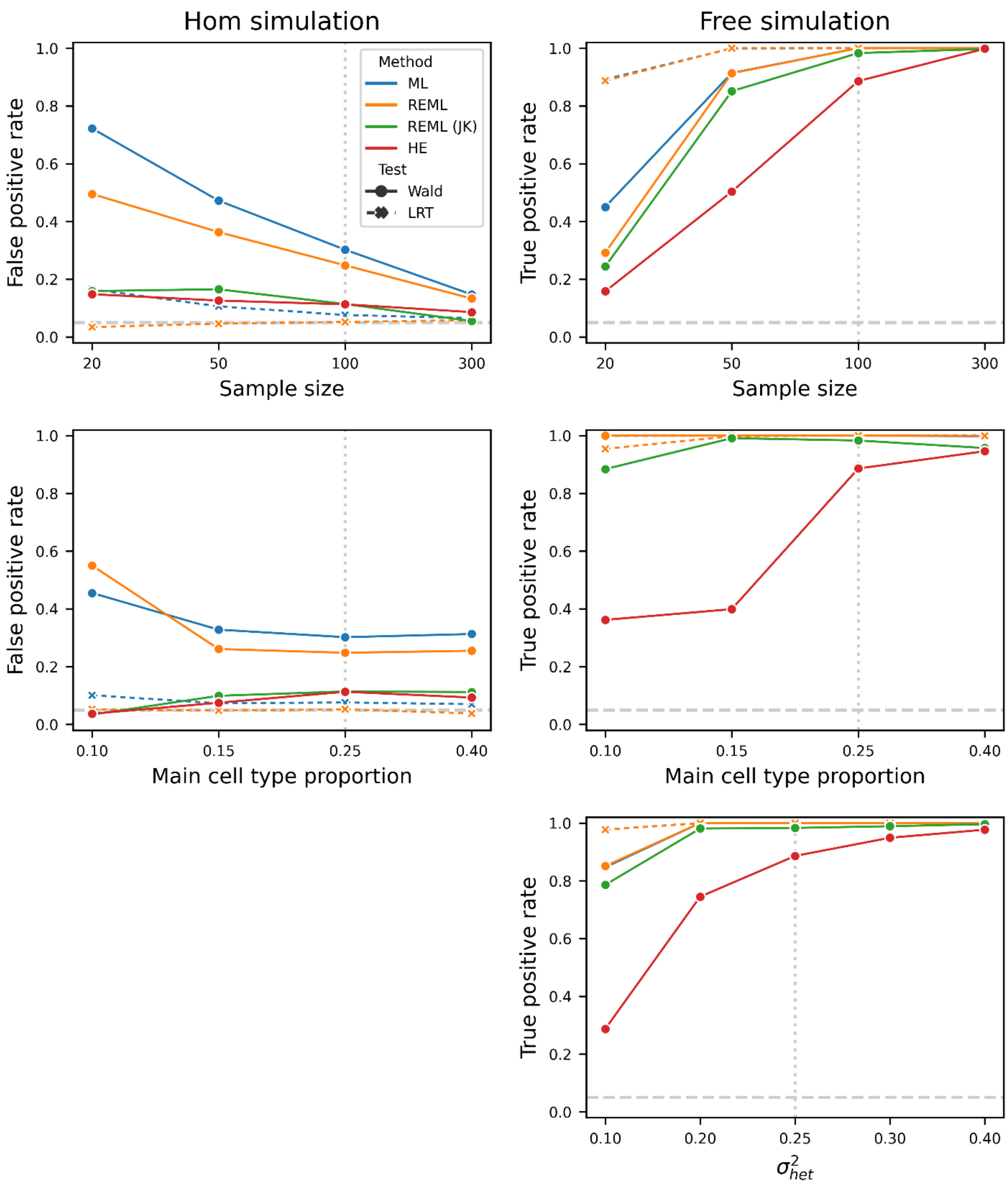


**Figure S6. Power of CTMM's test of cell type-specific variance with CTP data.** The left column shows simulations under the null Hom model, where there is no cell type-specific variance. The right column shows simulation under the alternative Free model, where each cell type has its own cell type-specific variance. Vertical dashed lines indicate parameters used in the baseline model as listed in Table S1. The top row varies sample size (as in Figure S1); the middle row varies cell type proportion (as in Figure S2); and the bottom row varies the overall level of cell type-specific variance ($\sigma^2_{het}$, as in Figure S3).

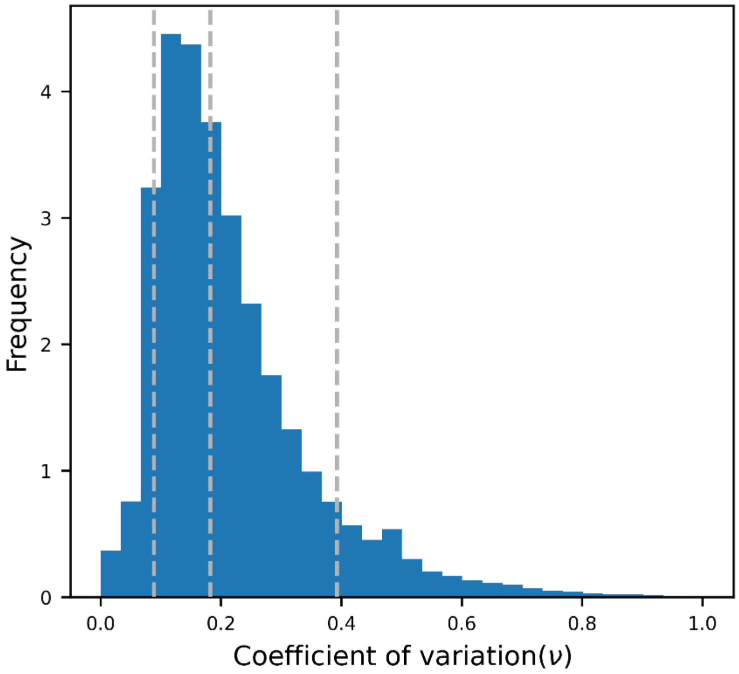


**Figure S7. Distribution of coefficient of variation for $\nu_{ic}$ across all combinations of individuals, cell types, and genes.** Dashed lines indicate the 10%, 50%, and 90% percentiles.

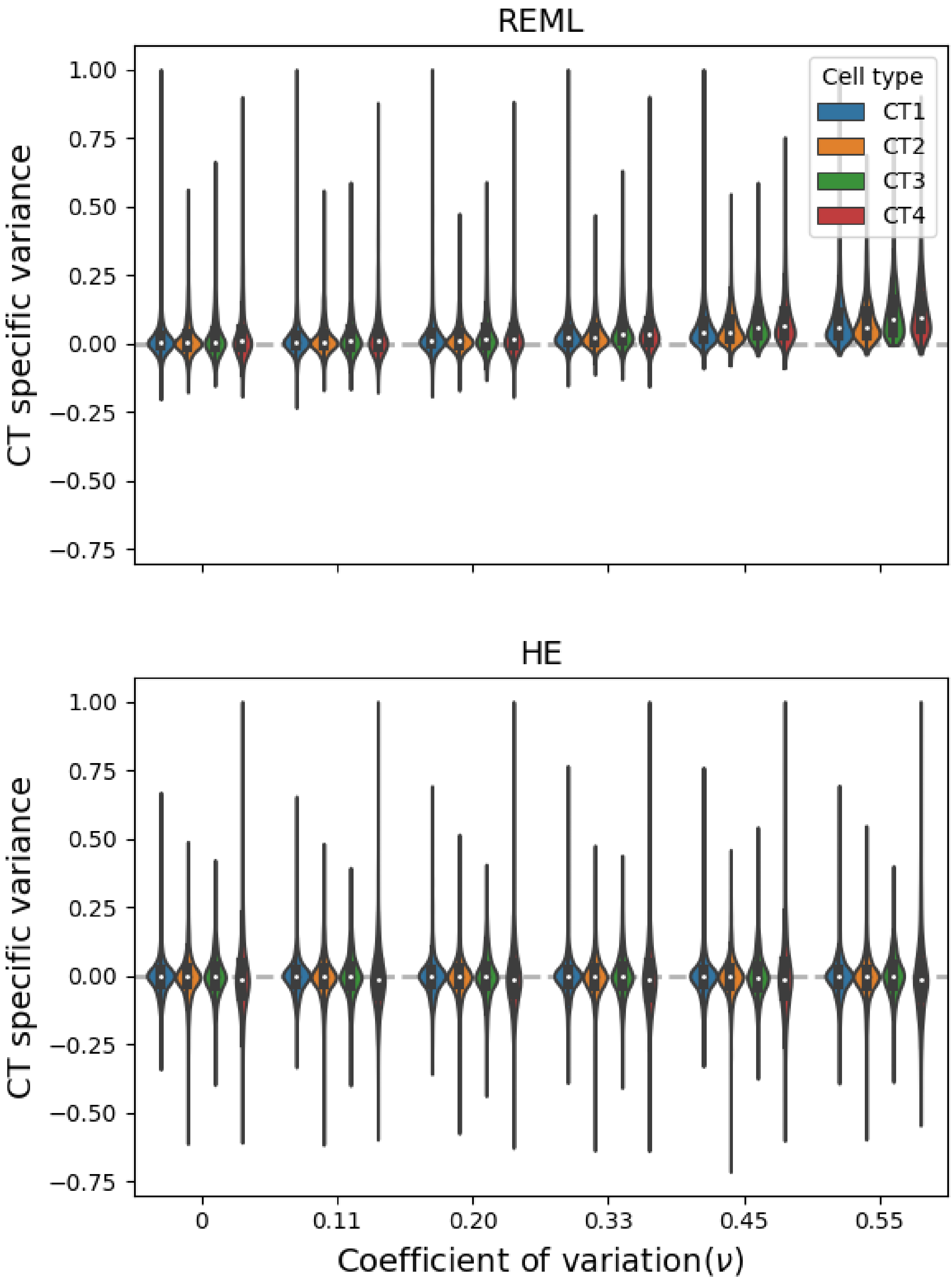


**Figure S8. CTMM estimates of cell type-specific variance in simulations of Hom model with noisy estimates of measurement error variance ($\nu_{ic}$).** Values above 1 were truncated.

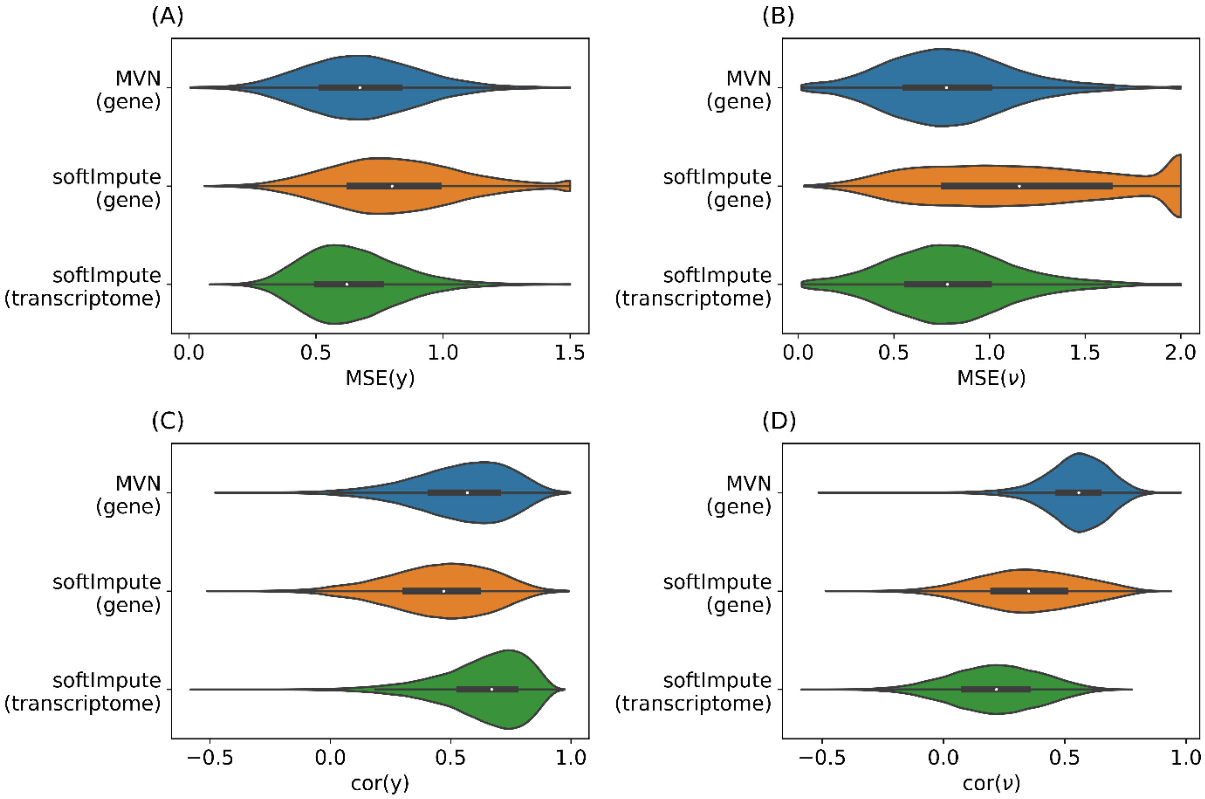


**Figure S9. Estimated imputation accuracies for cell type-specific pseudo-bulk and noise variance.** Mean squared error (MSE) (A, B) and correlation (C, D) between raw and imputed cell type-specific pseudo-bulk (A, C) and noise variance (B, D). Each violin shows the distribution across the transcriptome. SoftImpute and MVN are two matrix imputation methods. "Gene" refers to imputing each gene separately; "transcriptome" refers to jointly imputing all genes. $MSE(y)$ was truncated at 1.5; $MSE(\nu)$ was truncated at 2.

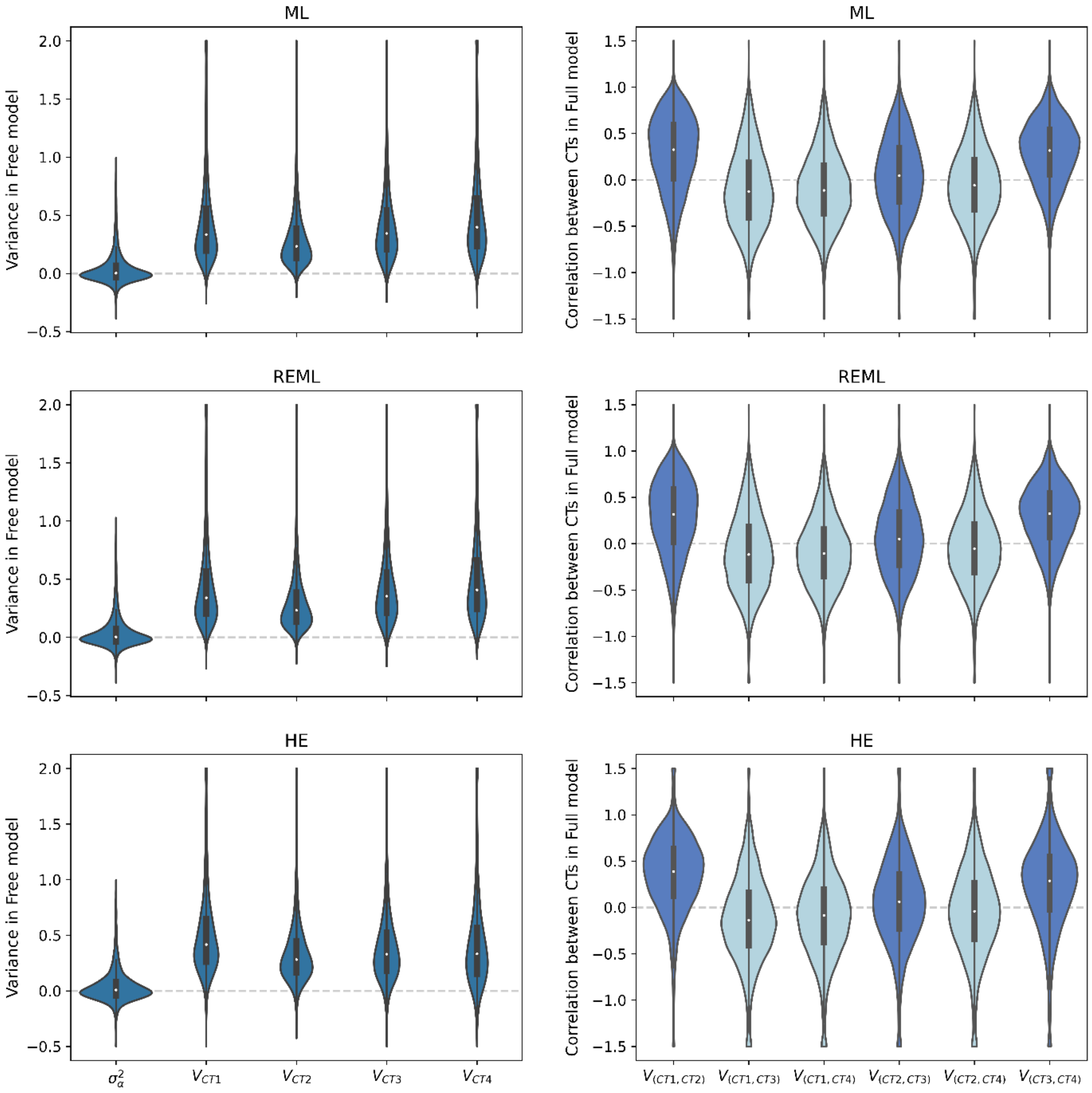


**Figure S10. Transcriptome-wide distribution of CTMM results with three estimation methods and CTP data.** Left column shows results from the Free model: homogeneous variance ($\sigma_\alpha^2$) and cell type-specific variances. Right column shows results from the Full model: correlation between each pair of cell types, with dark blue indicating adjacent cell type pairs and light blue indicating others. Homogeneous variance and cell type-specific variances were truncated to (-0.5,2); correlation between cell types was truncated to (-1.5,1.5).

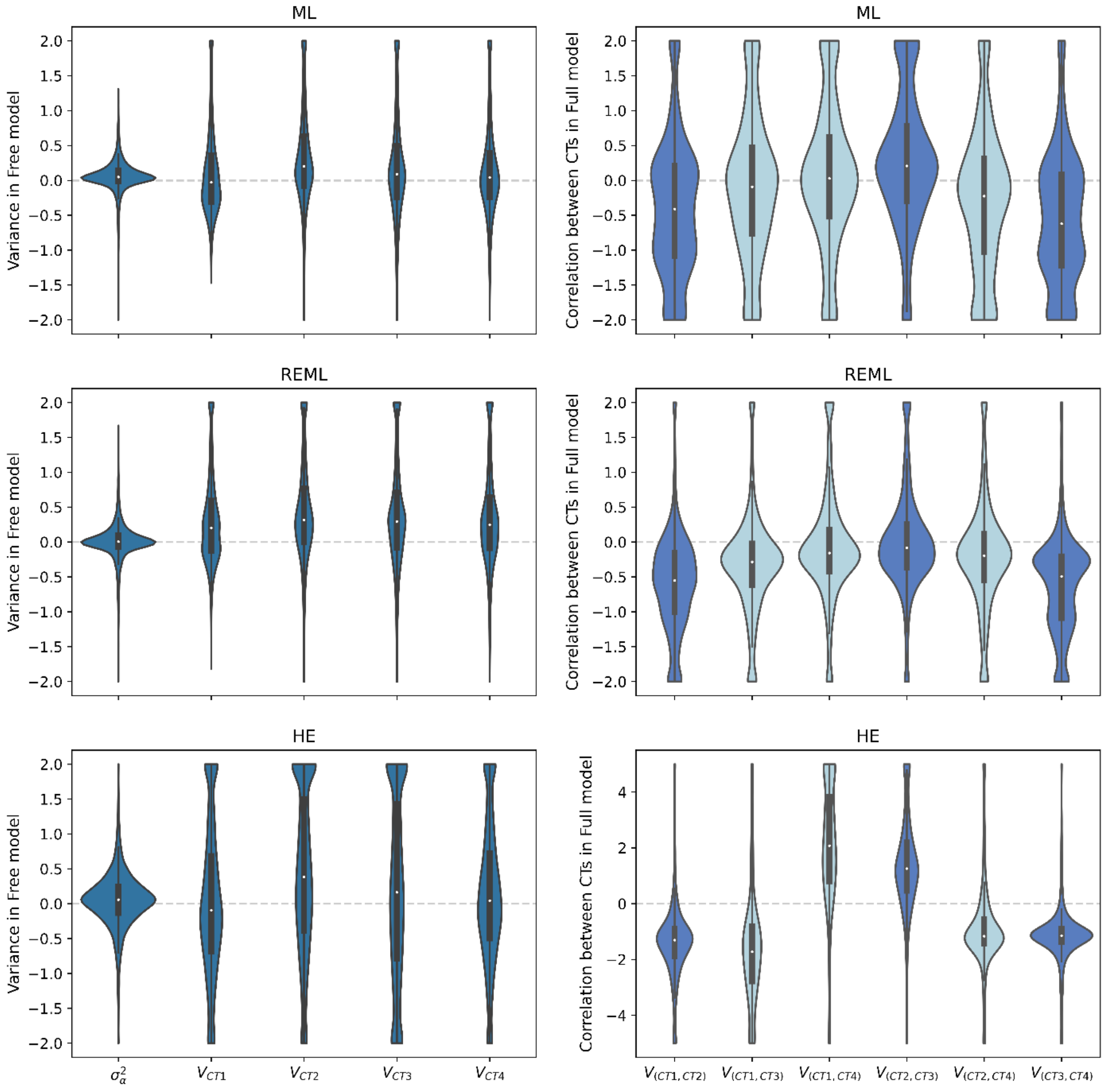


**Figure S11. Transcriptome-wide distribution of CTMM results with three estimation methods and OP data.** Left column shows results from the Free model: homogeneous variance ($\sigma_\alpha^2$) and cell type-specific variances. Right column shows results from the Full model: correlation between each pair of cell types, with dark blue indicating adjacent cell type pairs and light blue indicating others. Values were truncated to (-2, 2) except for the very noisy bottom-right panel which was truncated to (-5, 5).

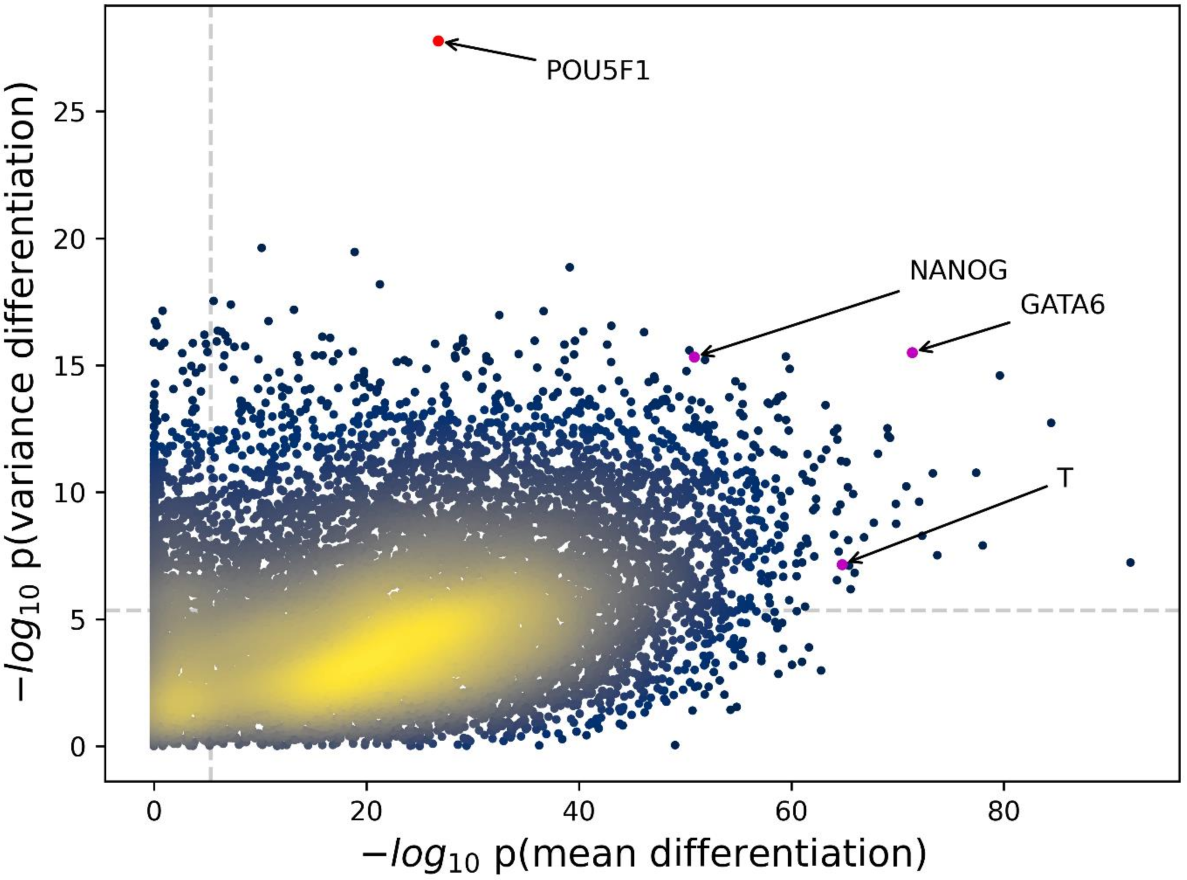


**Figure S12. Transcriptome-wide distribution of *p* values for differentiation in expression mean and variance using HE and CTP data.** Each dot represents a gene. Dots are colored by the density of genes in the area, with yellow indicating denser distribution. Dashed lines indicate significance threshold after Bonferroni correction. The three purple dots indicate the three maker genes used in *Cuomo et al*. to indicate each stem cell differentiation stage, spanning iPSC (*NANOG*), mesendoderm (*T*), and definitive endoderm (*GATA6*). The red dot indicates the top signal *POU5F1* found here and in REML with jackknife, which is one of the three core regulators in cell pluripotency.

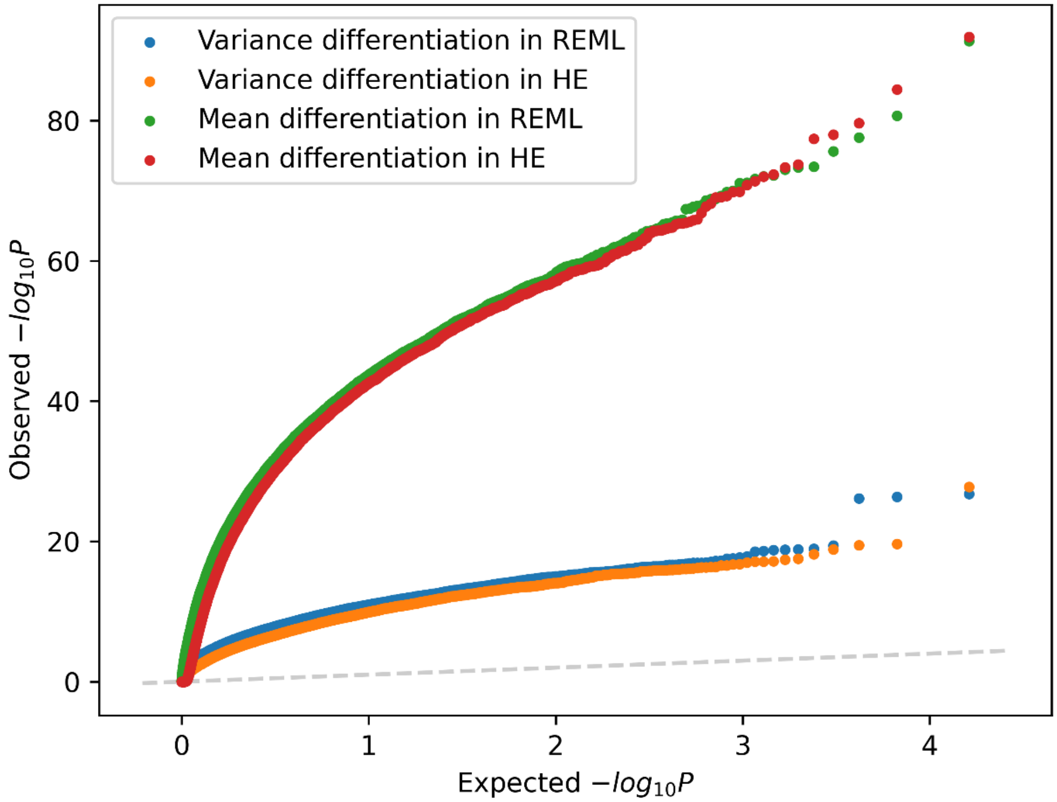


**Figure S13. Quantile-quantile plot of $p$-value distributions of variance differentiation and mean differentiation in REML and HE.**

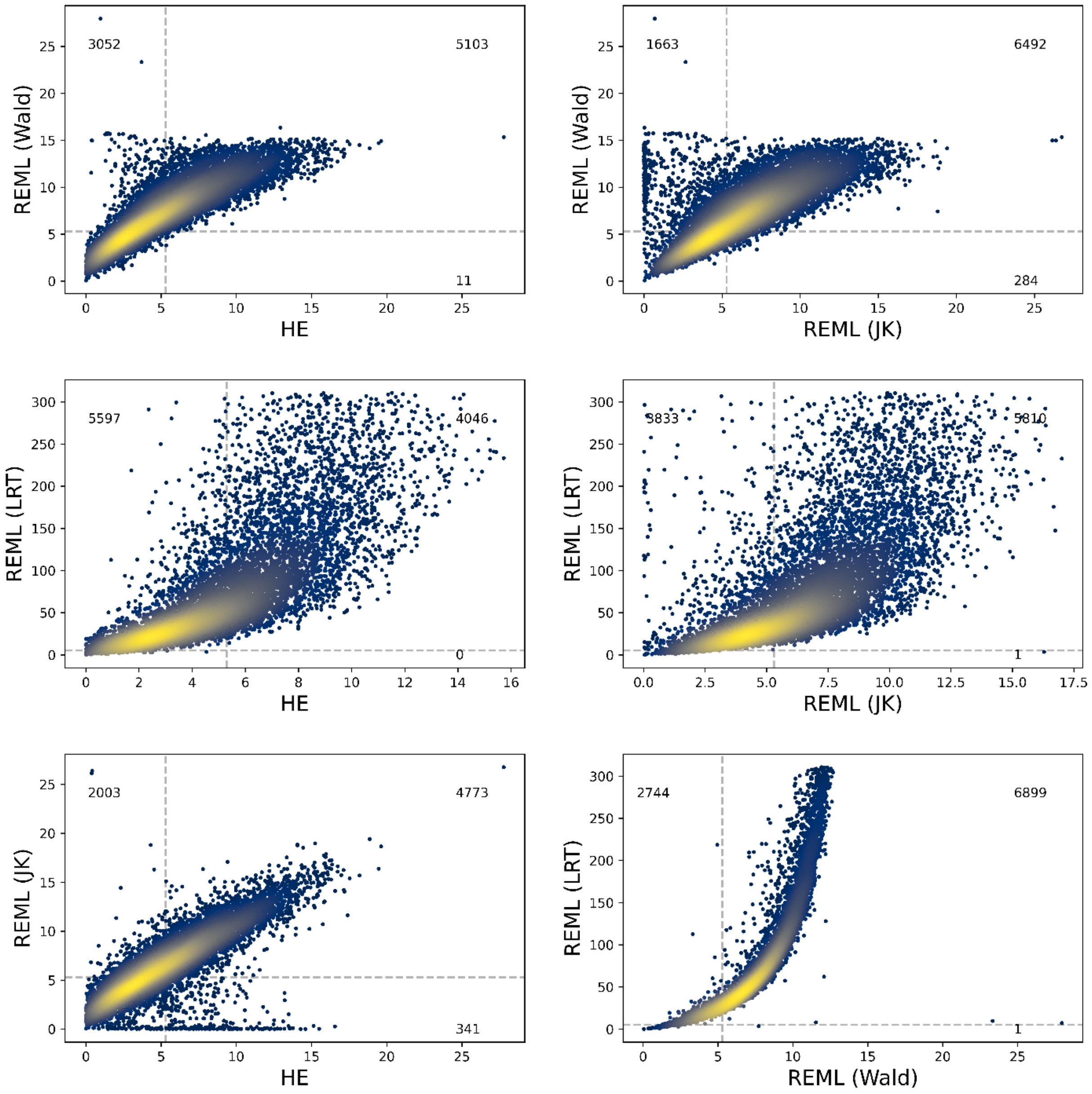


**Figure S14. Comparison of different tests for cell type-specific variance in REML and HE with CTP data.** Each axis shows $-\log_{10} p(\text{variance differentiation})$. Dashed lines indicate Bonferroni-adjusted significance thresholds ($p = 0.05/number\ of\ genes$). Wald tests require the precision matrix for parameter estimates, which is based on either the inverse of Fisher information matrix (for REML (Wald)) or jackknife (for REML (JK) and HE). Likelihood ratio tests (LRT) are also shown for REML.

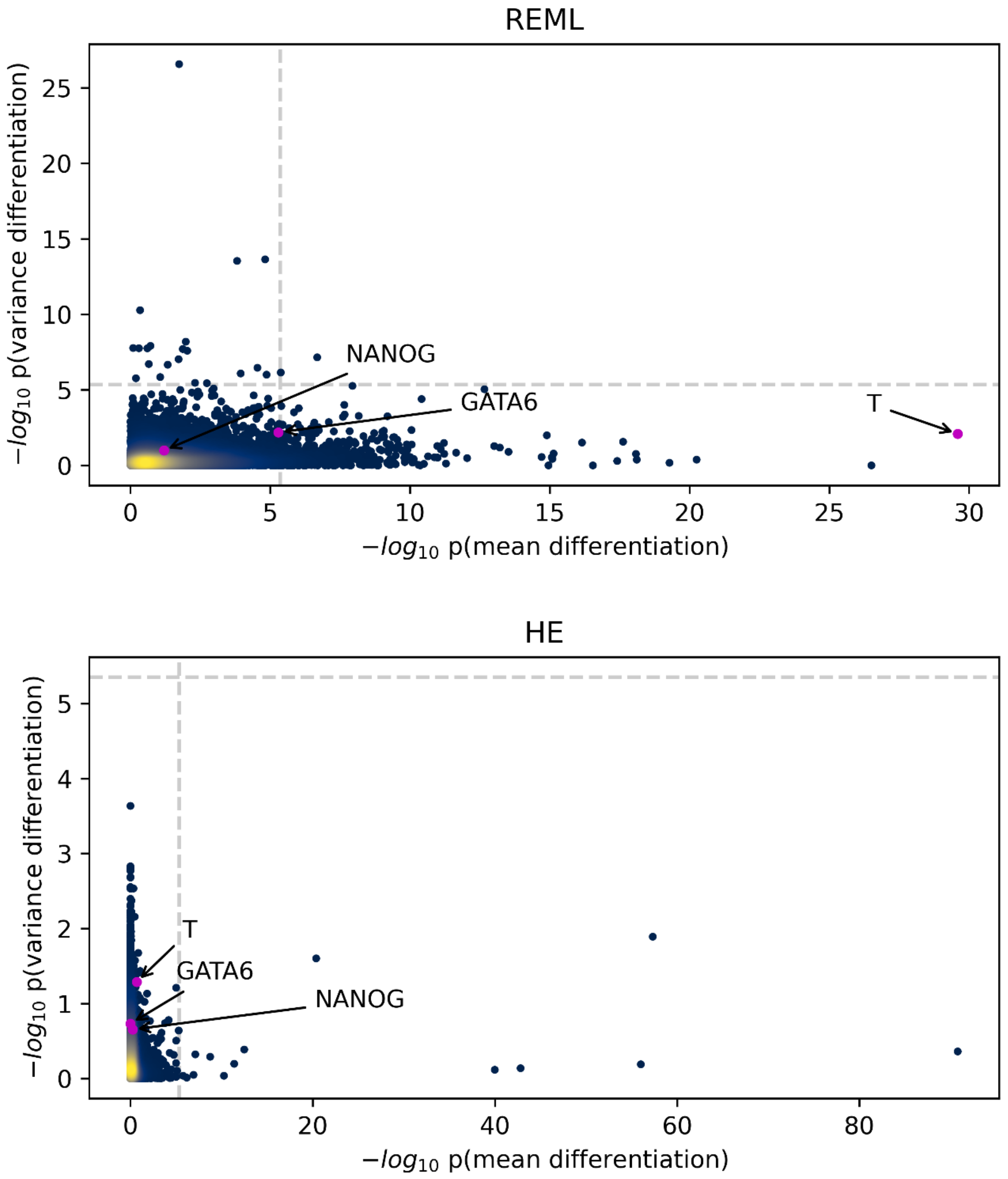


**Figure S15. Transcriptome-wide distribution of *p* values for differentiation in expression mean and variance using REML and HE with OP data.** In REML, $p$ values from Wald test were shown for mean differentiation, $p$ values from likelihood ratio test (LRT) were shown for variance differentiation. In HE, $p$ values for mean differentiation and variance differentiation were both from the Wald test. Each dot represents a gene. Dots are colored by the density of genes in the area, with yellow indicating denser distribution. Dashed lines indicate significance threshold after Bonferroni correction. The three purple dots indicate the three maker genes used in *Cuomo et al*. to indicate each stem cell differentiation stage, spanning iPSC (*NANOG*), mesendoderm (*T*), and definitive endoderm (*GATA6*).
